# Dynamics of compartment-specific proteomic landscapes of hepatotoxic and cholestatic models of liver fibrosis

**DOI:** 10.1101/2024.03.26.586230

**Authors:** Marketa Jirouskova, Karel Harant, Pavel Cejnar, Srikant Ojha, Katerina Korelova, Lenka Sarnova, Eva Sticova, Christoph H. Mayr, Herbert B. Schiller, Martin Gregor

## Abstract

Accumulation of extracellular matrix (ECM) in liver fibrosis is associated with changes in protein abundance and composition depending upon etiology of the underlying liver disease. Current efforts to unravel etiology-specific mechanisms and pharmacological targets rely on several models of experimental fibrosis. Here, we characterize and compare dynamics of hepatic proteome remodeling during fibrosis development and spontaneous healing in experimental models of hepatotoxic (carbon tetrachloride (CCl_4_) intoxication) and cholestatic (3,5-diethoxycarbonyl-1,4-dihydrocollidine (DDC) feeding) injury. Using detergent-based tissue extraction and mass spectrometry, we identified compartment-specific changes in the liver proteome with detailed attention to ECM composition and changes in protein solubility. Our analysis revealed distinct time-resolved CCl_4_ and DDC signatures, with identified signaling pathways suggesting limited healing and a potential for carcinogenesis associated with cholestasis. Correlation of protein abundance profiles with fibrous deposits revealed extracellular chaperone clusterin with implicated role in fibrosis resolution. Dynamics of clusterin expression was validated in the context of human liver fibrosis. Atomic force microscopy of fibrotic livers complemented proteomics with profiles of disease-associated changes in local liver tissue mechanics. This study determined compartment-specific proteomic landscapes of liver fibrosis and delineated etiology-specific ECM components, providing thus a foundation for future antifibrotic therapies.

## INTRODUCTION

Liver fibrosis and subsequent cirrhosis, two leading causes of liver disease-related deaths worldwide,^1^ develop as a result of chronic liver injury of multiple etiologies.^2^ Fibrosis is characterized by excessive accumulation of extracellular matrix (ECM) proteins forming fibrous scar tissue. In the liver, such pathological matrix is deposited by activated hepatic stellate cells (HSCs) and/or portal fibroblasts (PFs) in response to the inflammatory reaction.^2, 3^ Recent multi-omics studies have framed fibrosis as a dynamic multicellular process associated with the remodeling of gene expression landscapes^4^ and profound changes in liver protein abundance as well as composition.^5-7^

Hepatocellular injury alters parenchymal mechanical properties,^8^ thus further facilitating activation of resident HSCs and PFs and their differentiation into myofibroblasts depositing collagen-rich ECM.^9^ Accumulation of abnormal ECM further promotes ECM-depositing myofibroblasts and fosters progressive whole-organ stiffening as a result of ongoing fibrogenesis.^10^ Thus, ECM-associated changes are increasingly perceived as causative, rather than consequential, and multiple efforts have focused on the identification of extracellular niche components that drive the pathogenesis of fibrosis.^11^

The ECM is a complex network of hundreds of proteins acting as a three-dimensional scaffold, supporting cells mechanically and functioning as a reservoir for secreted factors (e.g., growth factors and cytokines).^12^ This so-called “matrisome” consists of “core matrisome” (the structural components of the ECM; i.e., collagens, glycoproteins, and proteoglycans) and “matrix-associated matrisome” (the ECM-interacting proteins).^13^ Transitions of extracellular factors between the insoluble matrisome and the soluble pools modulate their signaling capabilities and bioavailability.^11^ As fibrous ECM assemblies determine also tissue mechanics, matrisome constituents provide matrix-embedded cells with spatially distinct biochemical and biomechanical context.

In terms of etiology, chronic liver injury is inflicted either by hepatotoxic or cholestatic insult.^2^ In recent decades, several corresponding animal models of liver fibrosis have been established.^14^ Despite several translational limitations,^15^ these experimental models mimic fundamental aspects of human liver fibrosis.^14^ For instance, iterative carbon tetrachloride (CCl_4_) intoxication causes hepatocellular damage, HSC activation, and development of pericentral liver fibrosis that evolves into severe bridging fibrosis.^14^ In contrast, 3,5-diethoxycarbonyl-1,4-dihydrocollidine (DDC) feeding induces obstructive cholestasis, characterized by expansion of PFs and consecutive periportal fibrosis with typical portal–portal septa.^16^ As both human and animal studies have shown that liver fibrosis can be ameliorated by either targeting progression or promoting resolution,^15^ experimental models have become instrumental for identifying factors and mechanisms central to blocking fibrosis progression and promoting the reversal of advanced fibrosis.

In this comparative study, we employed CCl_4_- and DDC-based mouse models to describe proteomic changes during liver injury, fibrosis development, and repair while focusing on matrisome remodeling and associated alterations in tissue biomechanics. Our in-depth analysis of MS data obtained from total liver lysates and ECM-enriched insoluble fractions defines compartment- and time-resolved proteomic signatures in hepatotoxin-versus cholestasis-induced fibrosis and healing and delineates disease-specific matrisome.

## MATERIALS AND METHODS

### Animals and liver injury models

C57BL/6J male mice 8–10 weeks old were used for the analysis. All experiments were performed in accordance with an animal protocol approved by the Animal Care Committee of The Institute of Molecular Genetics and according to the EU Directive 2010/63/EU for animal experiments. To induce cholestatic liver injury, mice were either injected with 1 μl/g CCl_4_ ^14^ or fed a diet supplemented with 0.1% DDC.^16^

### Mass spectrometry sample preparation

A sample of ca 1 mm^3^ excised from the middle section of left lateral liver lobe was cryo-homogenized using TissueLyser II (QIAGEN, Germantown, MD, USA), resuspended, and sonicated. An aliquot of the sample was precipitated and labeled “Total”. The remaining solution was centrifuged at 16,000 × g for 20 minutes to separate the supernatant containing soluble (S)-fraction from the pellet, which itself contained insoluble ECM-enriched (E)-fraction.

### Nano-liquid chromatography mass spectrometry analysis

Nano reversed-phase columns were used for liquid chromatography MS analysis. Eluting peptide cations were analyzed on a Thermo Orbitrap Fusion collision cell, linear ion trap mass analyzer (Q-OT-qIT, Thermo Scientific). Tandem MS was performed by isolation at 1.5 Th with the quadrupole, higher-energy collisional dissociation fragmentation, and rapid scan MS analysis in the ion trap. Peptides were identified and quantified by MaxQuant label-free quantification software (1.6.7 version) using *Mus musculus* UniProt protein database (UniProtKB version July 2020).

### Human samples

The use of completely anonymized archived liver tissue samples for research purposes has been approved by the Ethical Committee of the Institute of Experimental and Clinical Medicine and Thomayer University Hospital, Prague, Czech Republic. Written informed consent was obtained from all patients enrolled in the study. All research was conducted in accordance with both the Declarations of Helsinki and Istanbul. Processing of human liver sections for immunofluorescence is detailed in the Supporting Information.

### Atomic force microscopy

Atomic force microscopy (AFM) was performed on liver sections 30 μm thick as previously described.^17^ Using the force mapping method, we measured 30 × 100 μm^2^ (10 × 36 pixels) defined regions precisely located by polarized microscopy. Seven areas (two sections per mouse) were chosen from three different mice for each time point (control, T1, T2, and T4).

### Statistics

All graphs and statistical tests indicated in graphs were performed using GraphPad Prism (GraphPad Software). All results are presented as mean ± SEM unless indicated otherwise. In the boxplots, the box represents 25th and 75th percentiles with the median indicated; whiskers reach to the last data point. Statistical tests used are specified in the figure legends. Statistical significance was determined at levels *, *p* < 0.05, **, *p* < 0.01, and ***, *p* < 0.001.

For further details regarding the materials and methods, please refer to the Supporting Information.

## RESULTS

### Proteomic analysis outlines the gradual fibrosis development and partial healing in both CCl_4_- and DDC-induced liver injury

To compare time-resolved, compartment-specific proteomes of CCl_4_- and DDC-induced models of liver fibrosis, total liver lysates (Total) together with two protein fractions obtained by one-step detergent extraction (ECM-enriched insoluble fraction (E-fraction) and soluble fraction (S-fraction)) were prepared for MS analysis (Figure 1A). The gradual fibrogenesis was assessed at two time points (T1 and T2) corresponding to mild fibrosis with partially bridging septa (T1, 3 weeks for CCl_4_ and 2 weeks for DDC treatment) and advanced fibrosis (T2, 6 weeks for CCl_4_ and 4 weeks for DDC). To characterize the dynamics of spontaneous fibrosis resolution, we allowed the mice 5 (T3) or 10 days (T4) to recover. Both models developed typical progressive liver damage with fibrotic scarring followed by partial healing upon insult withdrawal as demonstrated by plasma levels of liver injury markers and the extent of collagen-rich deposits in liver sections (Figure S1).

**FIGURE 1.**
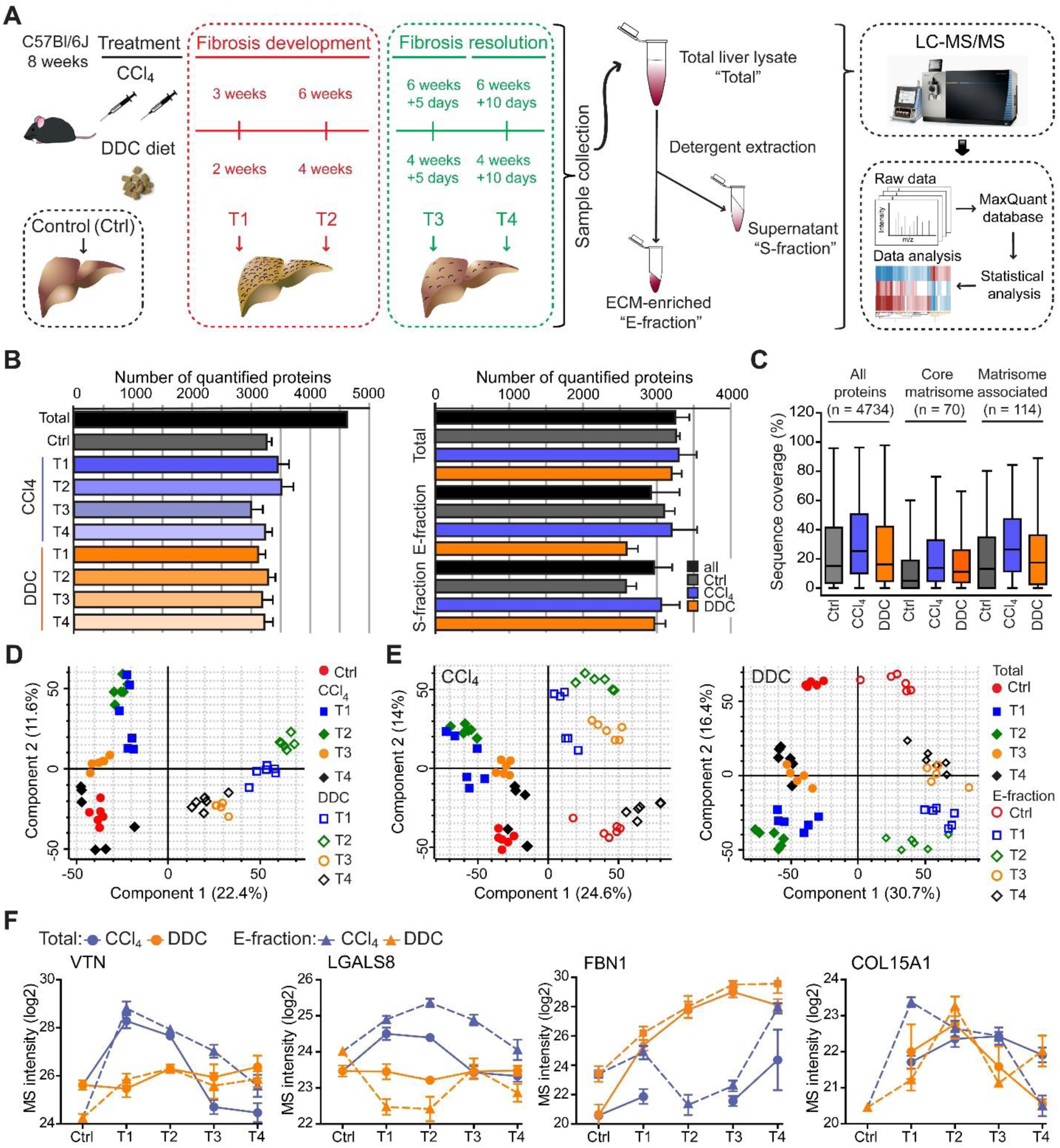
Hepatotoxic and cholestatic injury generate distinct time-resolved and compartment-specific protein signatures. (**A**) Schematic overview of the experimental setup. Six animals were used in each model at each time point. (**B**) Numbers of quantified proteins at the indicated time points and experimental conditions; n = 6. (**C**) Box plot shows the distribution of protein sequence coverage (coverage of possible tryptic peptides per protein in %) for the indicated matrisome categories (as defined by Naba et al.^13^) and all detected proteins in experimental conditions indicated. (**D**) Principal component analysis (PCA) of Total proteome separates time-dependent fibrogenesis and healing in CCl_4_-(closed symbols) and DDC-(open symbols) induced fibrosis. The first two components of data variability of 3,624 proteins identified in Total liver fractions in CCl_4_ and 3,521 proteins in DDC are shown; n = 6. (**E**) PCA shows the separation of Total (closed symbols) and E-fraction (open symbols) proteomes in time; the first two components of data variability are shown; n = 6. (**F**) Line plots show time-dependent changes in mass spectrometry intensities in Total (solid line) and E-fraction (broken line) proteomes for indicated selected proteins in CCl_4_ and DDC models; n = 4–6.

Using the two models, we quantified 4,737 proteins approximately evenly distributed across all time points and fractions (Figure 1B, C, Supporting Results). Principal component analysis (PCA) revealed clear temporal separation of CCl_4_- and DDC-derived samples in both Total and E-fraction (Figure 1D, E). Unsupervised hierarchical cluster analysis of total lysate proteins together with annotation enrichment of the observed clusters further demonstrated substantial differences in the dynamic regulation of protein expression between the two models (Figure S2). This was evidenced by differential Total MS protein intensity temporal profiles of matrisome-enriched cluster constituents, such as vitronectin, galectin 8, fibrillin 1, or a putative portal myofibroblast marker collagen α-1(XV) chain (Figure 1F). Moreover, distinct galectin 8 and vitronectin expression profiles obtained for CCl_4_ and DDC E-fractions revealed complex changes in protein association with the insoluble proteome. Taken together, our MS data allow for time-resolved identification of proteins differentially expressed during fibrosis progression and resolution in etiologically distinct models of liver fibrosis. Moreover, this approach enables analysis of complex changes in the solubility of the identified proteins and their transition between liver compartments.

### Time-resolved analysis of Total proteomes indicates limited healing and potential tumorigenicity in the DDC model

To capture the proteome differences between hepatotoxin-vs. cholestasis-induced fibrogenesis, we determined proteins significantly regulated during fibrogenesis separately in CCl_4_ and DDC Total samples (Benjamini-Hochberg false discovery rate (B.H. FDR) < 5%; see Supporting Materials and Methods). In total, 702 proteins were found to be shared by the two models, while 514 (CCl_4_) and 1,074 (DDC) proteins were identified as unique for the respective model (Figure 2A).

**FIGURE 2.**
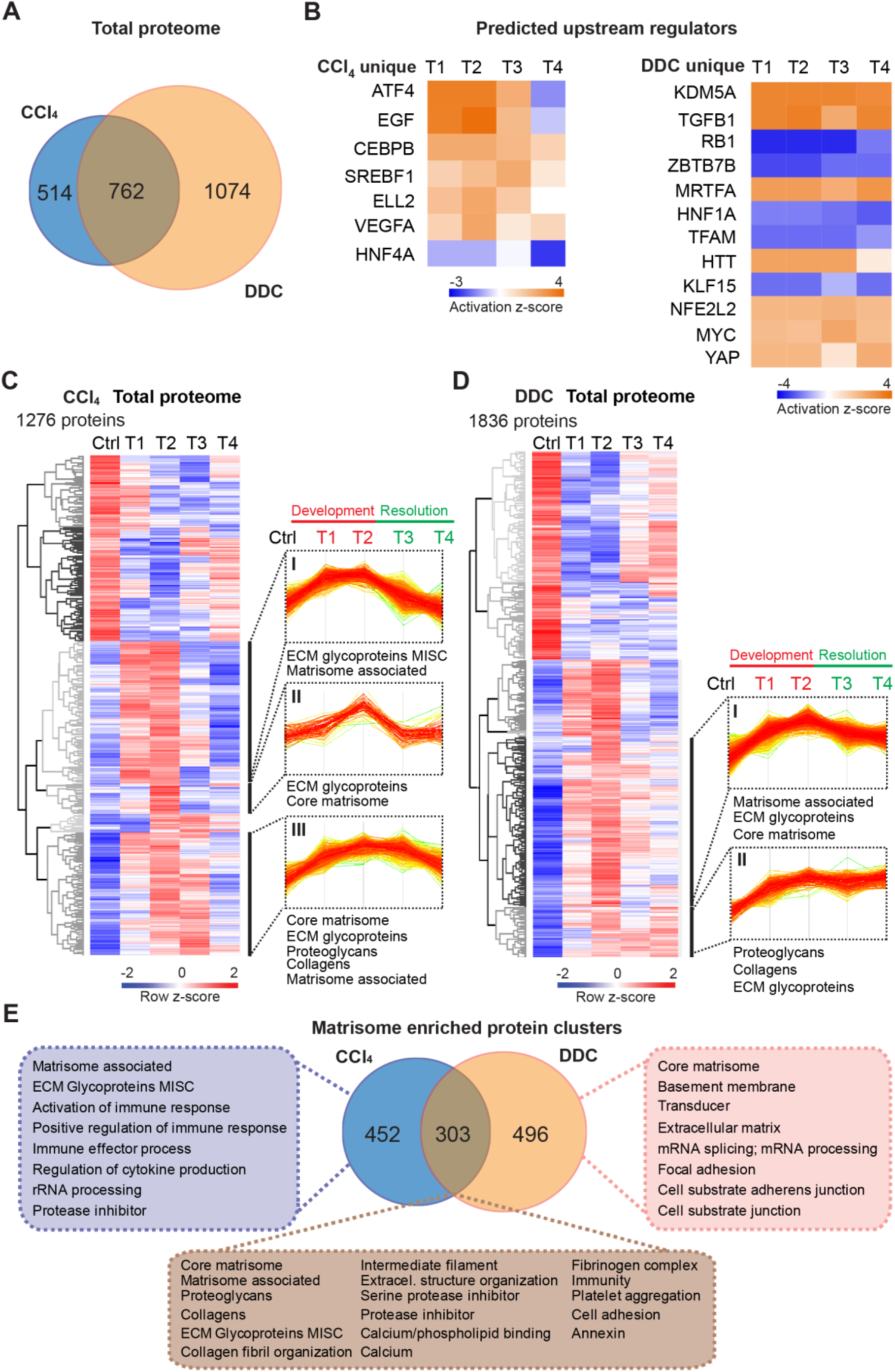
Time-resolved analysis of Total proteomes shows limited healing in matrisome-enriched protein clusters in the cholestatic model. (**A**) Venn diagram shows relative proportion of 1,276 and 1,836 proteins differentially expressed (*t*-test; Benjamini-Hochberg FDR < 5%) in Total CCl_4_ and DDC proteomes. (**B)** Hierarchical cluster analysis of the activity score of the upstream regulators at the indicated time points predicted by Ingenuity Pathway Analysis (IPA) from unique CCl_4_ and DDC protein signatures shown in A. (**C,D**) Hierarchical clustering of mean *z*-scored mass spectrometry (MS) intensities of proteins of Total CCl_4_ (C) or DDC (D) proteomes; n = 6. Profiles of *z*-scored MS intensities of proteins from matrisome-annotated clusters for CCl_4_ (I–III) and DDC (I and II) models are shown. (**E**) Venn diagram compares proteins from matrisome-annotated clusters shown in C and D. UniProt keyword enrichment annotation for each group within the diagram is indicated (Fisher’s test, Benjamini-Hochberg FDR < 4%).

Using Ingenuity Pathway Analysis (IPA), we predicted upstream transcriptional regulators and growth factors together with corresponding downstream biological signaling pathways differentially associated with identified protein groups (Figures 2B and S3A). Abundance changes in known targets indicated the activity of key regulators involved in fibrogenesis (TGFβ1 and angiotensinogen), hypoxia response (HIF1A), and inhibition of factors mediating hepatocyte function (HNF4α and HNF1α) or involved in HSC inactivation (PPARGC1A) in both fibrotic models (Figure S3A). Interestingly, the pro-proliferative mTOR pathway showed signaling attenuation in the course of CCl_4_ treatment but upregulation in the case of DDC treatment, suggesting hyperplastic potential of the DDC model (Figure S3A). Consistently, analysis of the DDC signature revealed inhibition of tumor suppressor gene *Rb1* with simultaneous activation of potential oncogenic regulators KDM5A, MRTFA, MYC, and YAP, thus underscoring a potential carcinogenic risk of cholestatic model (Figures 2B and S3B). By contrast, the CCl_4_ signature indicated activation of EGF and VEGFα with simultaneous downregulation of the HNF4α (Figure 2B) supporting thus previous reports on the importance of these factors during hepatotoxic injury.^18-20^ While dynamic changes of regulatory networks and signaling pathways correlated well with fibrosis progression in both models, ineffective downregulation of profibrotic signaling in the DDC-induced model suggested a limited capacity of the liver to heal from cholestatic injury (Figures 2B and S3B).

Next, we analyzed CCl_4_ and DDC Total proteomes using unsupervised hierarchical clustering and functional annotation term enrichment (Figure 2C,D). This allowed the separation of proteins with similar temporal abundance profiles while simultaneously revealing clusters comprising matrisome-annotated proteins (see Materials and Methods). In the CCl_4_ proteome, three matrisome-annotated clusters (Figure 2C; clusters I, II, and III) containing 755 proteins showed a variable degree of time-dependent abundance decline over the course of healing. By contrast, two large matrisome-annotated clusters (Figure 2D; clusters I and II) with 799 constituents showed almost no decline in abundance over the recovery period in the DDC model. Our data reflect the reversibility of CCl_4_-induced fibrogenesis and restricted capacity to heal with higher potential of carcinogenic risk in the DDC model.

### Matrisome linked with cholestasis is enriched in basement membrane (BM) proteins whereas deposits upon hepatotoxic injury contain matrisome-associated proteins

To further dissect disease-specific processes associated with matrisome-enriched clusters (Figure 2C,D), we followed the functional annotation of proteins using a comprehensive annotation matrix compiled from Gene Ontology terms, UniProtKB keywords, and the MatrisomeDB matrisome database.^13, 21^ This revealed that 303 proteins shared by CCl_4_ and DDC Total proteomes were annotated not only with matrisome-related keywords but also as “Cell adhesion”, “Fibrinogen complex”, “Platelet aggregation”, and “Intermediate filaments”, thus indicating engagement in cellular interaction with ECM (Figure 2E). Consistently, top canonical IPA pathways included “Signaling by Rho family GTPases”, “Integrin signaling” and “Actin cytoskeleton signaling” (Figure S3C). As anticipated, 452 proteins uniquely identified in CCl_4_ matrisome-enriched clusters were also associated with inflammatory response, a well-described feature of this model.^22^ A group of 496 proteins exclusive for DDC was found to associate with “Basement membrane” and “Cell substrate junction” reflecting an increased abundance of BM components synthesized during the development of periductal fibrosis (e.g., laminins, α-chains of collagens IV, VI, and XVIII; Figure 2E).

In a group of 60 matrisome proteins significantly changing in both Total CCl_4_ and DDC proteomes (Figure 3A) there were mainly core matrisome proteins, such as collagens (e.g., α-chains of collagens I and V) and ECM glycoproteins (e.g., fibronectin, EMILIN1, and vitronectin). In contrast to previous reports,^7^ we found collagen VI, a protein regulating ECM contractility,^23^ upregulated during fibrogenesis in both models (Figure S4). In addition, we identified several matrisome-associated proteins (e.g., NGLY1, PLOD3, S100A4, MUG2, SERPINA7, SERPINF1, and P4HA1) not previously reported in liver fibrosis as uniquely induced in the CCl_4_ model.^21^ Among these, ECM glycoprotein lactadherin (MFGE8), has been shown to reduce liver fibrosis in mice.^24^ In contrast, BM matrisome proteins, such as laminins, collagens IV and XVIII, and perlecan (HSPG2) were exclusively upregulated in the DDC proteome (Figures 3A and S4).

**FIGURE 3.**
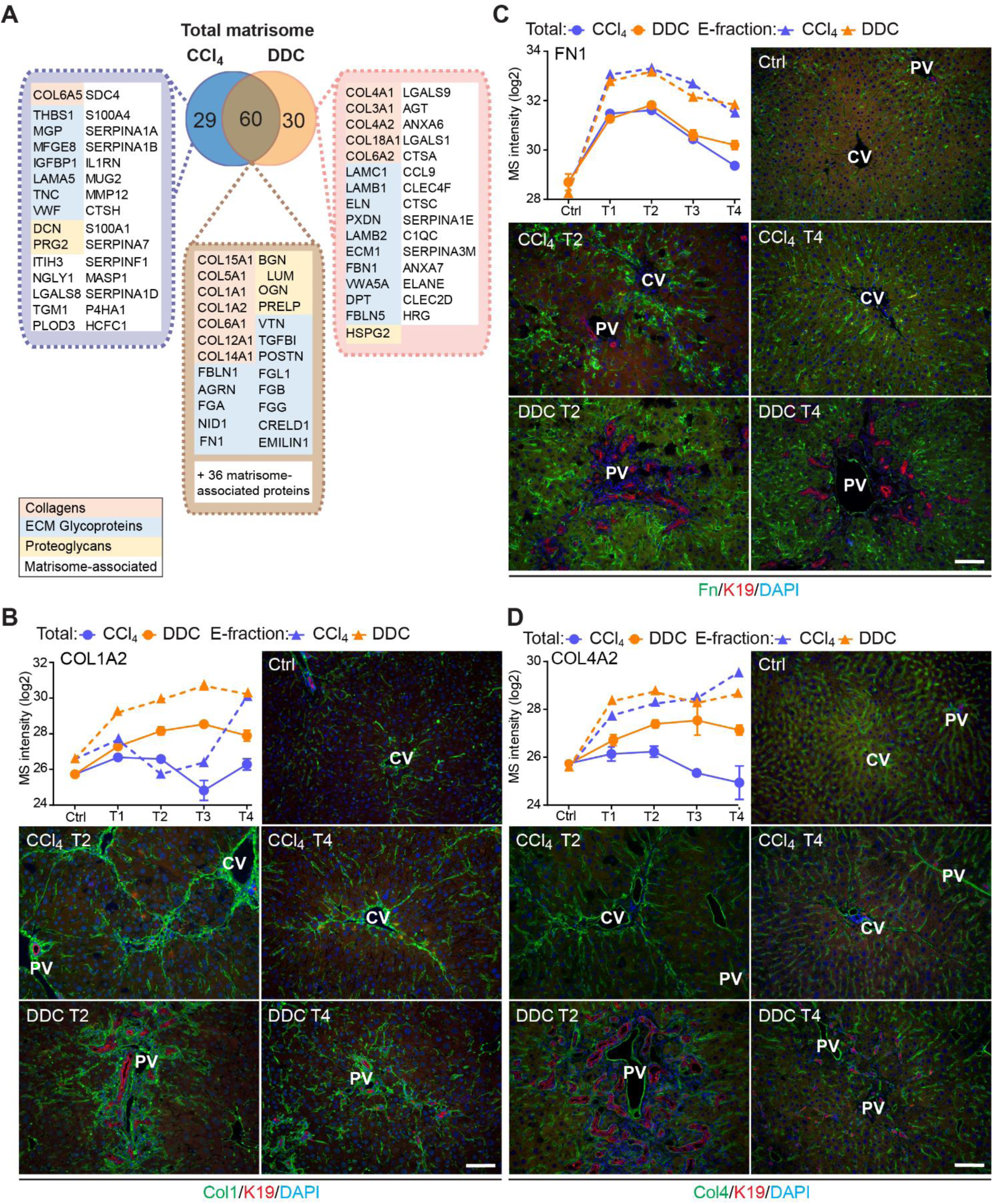
Comparison of matrisome proteins differentially expressed between the CCl_4_- and DDC-derived Total proteomes with immunofluorescence (IF) localization of the main core matrisome proteins within the injured livers. (**A**) Venn diagram shows a comparison of matrisome proteins differentially enriched in Total CCl_4_ and DDC proteomes. Color coding indicates identified matrisome categories. (**B–D**) Representative IF images of collagen I (B), fibronectin (C), and collagen IV (D), all in green in liver sections from untreated controls (Ctrl), CCl_4_-, and DDC-treated mice at time points of fibrosis development (T2) and resolution (T4). Bile ducts were visualized with antibodies to keratin 19 (K19; red); nuclei were stained with DAPI (blue). CV, central vein; PV, portal vein. Scale bar = 100 μm. Line plots show time-dependent change in respective mass spectrometry intensities in Total (solid line) and E-fraction (broken line) proteomes in CCl_4_ and DDC models; n = 4–6.

To validate our findings demonstrating distinct features of the fibrotic models used, we performed immunofluorescence (IF) analysis of major core matrisome proteins abundantly upregulated in both CCl_4_ and DDC proteomes (Figure 3B–D). Collagen I accumulated at the sites of primary injury and, consistent with proteomics, was only partially reduced upon recovery in the DDC model (Figure 3B). Collagen I-rich areas were delineated by fibronectin, a protein serving as a scaffold for collagen fibril assembly.^25^ Although fibronectin abundance in Total CCl_4_ and DDC proteomes decreased with healing, IF analysis revealed persisting deposits within capillarized hepatic sinusoids even after a recovery period of 10 days (Figure 3C). Consistent with a report on human cirrhotic livers,^26^ BM-associated collagen IV accumulated around fibrotic septa in both models and its persistent presence indicated incomplete liver healing after DDC withdrawal (Figure 3D).

### Integrin αv is specifically induced on the membranes of injured hepatocytes within fibrotic scar tissue in the hepatotoxic CCl_4_ model

The dynamic nature of fibrosis stems from interplay between injured hepatocytes, immune cells, and hepatic myofibroblasts.^3^ Using previously published cell type-specific signatures (see Materials and Methods), we identified 10 different cell populations in CCl_4_ and DDC samples and compared their abundances over the course of liver fibrosis (Figures 4A–D and S5A). A set of 18 proteins quantified from the signature of hepatocytes exhibited faster decline in the hepatotoxic CCl_4_ than DDC model (Figure 4A), corresponding well to the extensive parenchymal injury evidenced by high alanine and aspartate transaminase levels (Figure S1A). In support, large areas of injured pericentral hepatocytes were found negative for HNF4α staining in CCl_4_-model (Figure S6A). Rapid increase in HSC and activated PF signatures paralleled the activation and proliferation of ECM-depositing myofibroblasts in both fibrotic models (Figure 4B) evidenced by IF staining of myofibroblast marker αSMA (Figure S6A). However, faster upregulation of activated PFs in DDC confirms their predominant role in cholestatic fibrosis. Temporal abundance profiles of Kupffer cell, macrophage, monocyte, and B-cell signature proteins illustrate faster recovery from inflammation with healing in the CCl_4_ than in DDC model (Figures 4C,D and S5A). This was further confirmed by IF staining of liver sections (Figure S6B). These findings, together with matrisome-enriched protein cluster analysis (Figure 2E), emphasized the role of the inflammation component in the context of fibrosis and underscored the model-specific involvement of PFs in cholestatic fibrosis.

**FIGURE 4.**
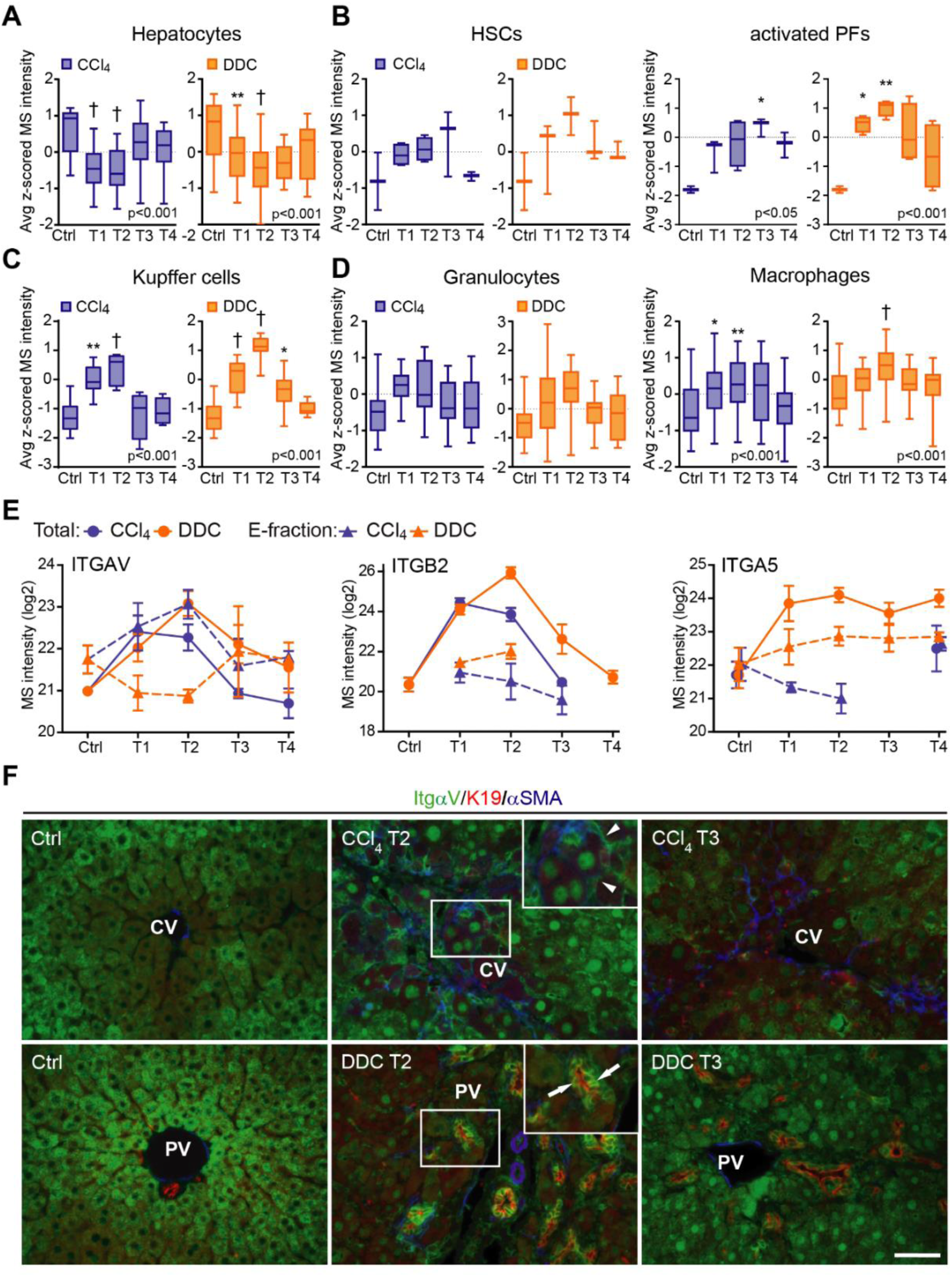
Liver cell type dynamics and cell type-specific integrin expression during fibrogenesis and healing. (**A–D**) Box plots show mean *z*-scored mass spectrometry (MS) intensities of the indicated cell type-specific protein signatures in time. Hepatocytes (A; n = 18), HSCs, and activated portal fibroblasts (PFs) (B; n = 3 and 4), Kupffer cells (C; n = 6), granulocytes, and macrophages (D; n = 16 and 33). One-way ANOVA with Bonferroni’s post-test; \**p* < 0.05; \*\**p* < 0.01; †*p* < 0.001. (**E**) The line plots show the time-dependent change in MS intensities of indicated selected integrins in Total (solid line) and E-fraction (broken line) proteomes in CCl_4_ and DDC models; n = 4–6. (**F**) Representative immunofluorescence images of liver sections from untreated controls (Ctrl), CCl_4_-, and DDC-treated mice at time points of fibrosis development (T2) and resolution (T4) immunolabeled for integrin αv (green), K19 (red), and αSMA (blue). Arrowheads, integrin αv-positive injured hepatocytes; arrows, integrin αv-positive biliary epithelial cells of reactive ductuli. CV, central vein; PV, portal vein. Boxed areas, ×2 images. Scale bar = 50 μm.

Cellular interactions with the changing microenvironment are mediated by integrins, a heterogeneous family of cell adhesion receptors. Thus, temporal changes in cell type-specific integrin subsets reflect the alterations in fibrotic injury-induced cell populations with the potential to predict the fibrogenicity of the ongoing disease. Our profiling revealed increased expression of integrin β1 (Figure S5B), the most abundant collagen receptor, which has been reported (together with integrins α1 and α5) to correlate with the stages of human liver fibrosis.^27^ Given its cell type-specific expression prevalence, this finding likely corresponds to the expansion of HSCs, simultaneously with induced integrin β1 expression on LSECs and injured hepatocytes.^27^ A rapid immune response is documented by extensive upregulation of main leukocyte integrin subunits β2 and αM (Figures 4E and S5B). Their differential expression kinetics between the models correlate well with cell type-specific signatures (Figures 4C,D and S5A). Unexpectedly, fibronectin receptor integrin α5 (typically expressed on HSCs, PFs, and LSECs^28, 29^ was detected in Total DDC proteome only (Figure 4E). Inasmuch as a boosted expression of integrin α5 has been reported in liver tumors,^30^ this finding further supports the tumorigenic potential of biliary fibrosis.

Strong induction of TGFβ-activating integrin αv in both CCl_4_ and DDC Total proteomes (Figure 4E) reflected its central role in fibrogenesis.^31^ Strikingly, in E-fraction, integrin αv was found to increase only in the CCl_4_ model (Figure 4E), suggesting its close association with ECM specifically during hepatotoxic injury. Subsequent IF analysis showed localization of antibodies to αv mostly to periportal hepatocytes and with little signal found in central hepatocytes of control livers (Figure 4F). During CCl_4_-induced injury, αv staining was enhanced at the periphery of pericentral hepatocytes adjacent to the collagen-rich scars, reflecting ongoing liver periportalization with the injury.^22^ The zonal distribution of αv was partially restored with fibrosis regression. In DDC-driven cholestasis, antibodies to αv stained strongly reactive biliary epithelial cells (BECs), the main drivers of fibrogenesis in biliary fibrosis (Figure 4F). Thus, our analysis reveals stage-specific induction of integrin αv on the surface of pericentral hepatocytes that has been unrecognized to date and suggests its potential as a marker of reactive cholangiocytes in cholestasis.

### Analysis of solubility profile dynamics of liver proteome reveals extracellular matrix protein-1 among matrisome proteins induced by fibrogenesis

As ECM remodeling during fibrogenesis entails also changes in the solubility of its constituents and associated proteins,^11^ we next set out to identify proteins that become increasingly insoluble with fibrosis progression. We found 1,273 (CCl_4_) and 762 (DDC) differentially expressed proteins, defined by at least 3-fold higher expression in E-than in S-fraction (see Materials and Methods). Hierarchical clustering of their MS intensity ratios clearly reflected changes in protein solubility profiles in the course of disease progression (Figure S7A,B). Proteins grouped into three (CCl_4_, 406 proteins) and two (DDC, 63 proteins) ECM-enriched clusters exhibited increasing insolubility during fibrogenesis. Matrisome proteins found in both fibrotic models were mainly collagens and ECM glycoproteins (Figure 5A). Interestingly, we identified 93 (CCl_4_) and 16 (DDC) proteins not present in control samples but induced by treatment (Figure S7C, Supporting Results). Abundance of most of these proteins in E-fraction decreased with healing in the CCl_4_ but not in the DDC model (Figure 5B), suggesting model-specific incorporation of matrisome-associated proteins into insoluble ECM.

**FIGURE 5.**
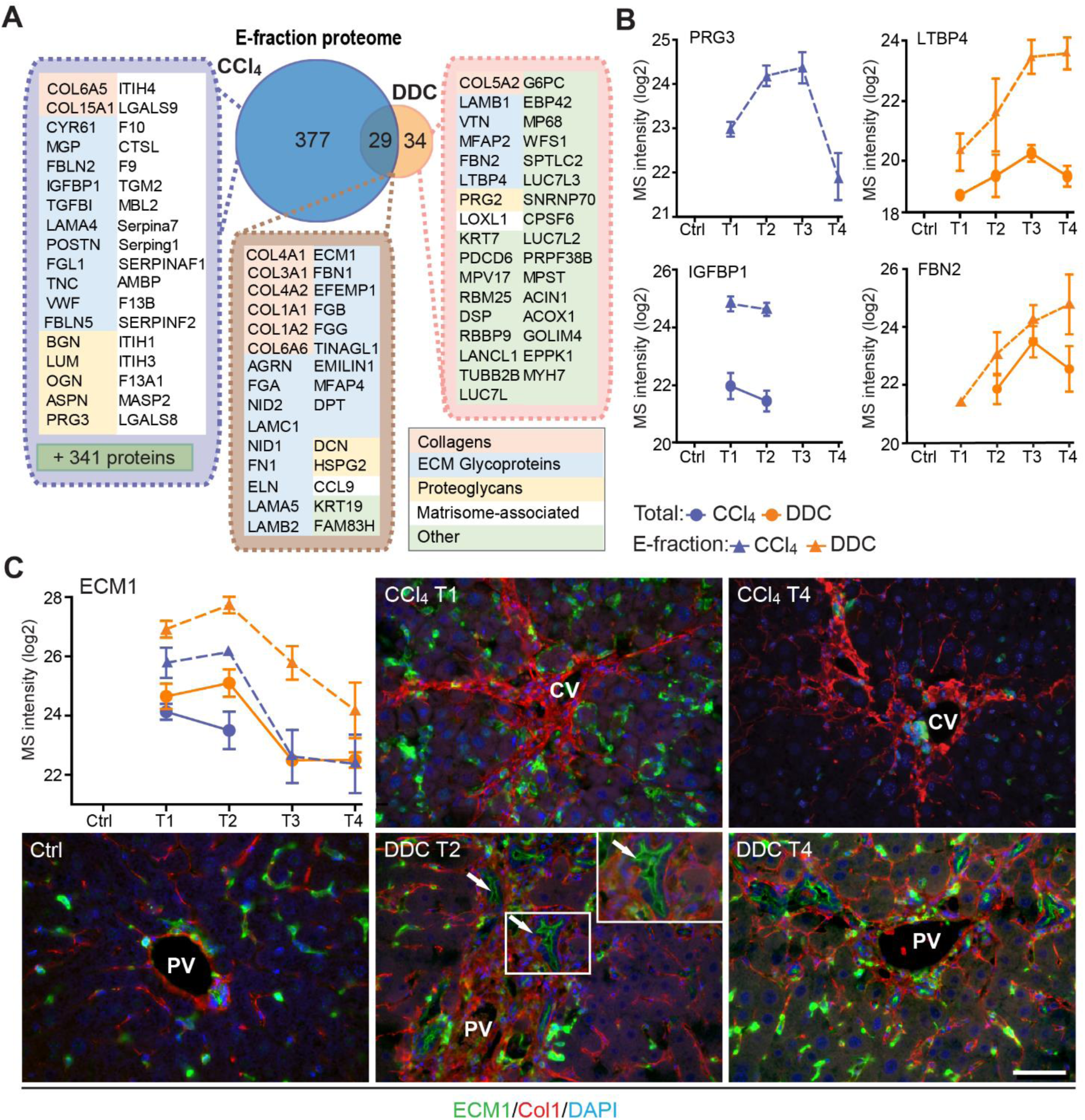
Solubility profiling provides in-depth analysis of model-specific matrisome composition. (**A**) Venn diagram shows relative proportion of proteins from E-fraction proteome identified as proteins with increasing insolubility over the course of fibrosis in each model (see Figure S7 and Supporting Materials and Methods). Matrisome proteins are highlighted with color coding to indicate identified matrisome categories. (**B**) Line plots show time-dependent change in mass spectrometry (MS) intensities of indicated selected matrisome proteins uniquely identified in E-fraction (broken line) proteomes in CCl_4_ and DDC models; n = 4–6. (Solid line shows MS intensity profile in Total proteome.) (**C**) Representative immunofluorescence images of liver sections from untreated controls (Ctrl), CCl_4_-, and DDC-treated mice at indicated time points of fibrosis development (T1 and T2) and resolution (T4) immunolabeled for ECM1 (green) and collagen 1 (red). Nuclei were stained with DAPI (blue). Arrows, ECM1-positive reactive biliary epithelial cells. CV, central vein; PV, portal vein. Boxed areas, ×2 images. Scale bar = 50 μm.

Among proteins upregulated with disease onset in E-fractions of both fibrotic models was extracellular matrix protein-1 (ECM1; Figures 5C and S7C). Subsequent IF analysis confirmed weak ECM1 staining (corresponding to low expression levels) in control livers, which was confined mostly to Kupffer cells and the apical membrane of BECs (Figures 5C and S7D). Increased ECM1 abundance during fibrogenesis was paralleled by its increasing localization within the infiltrating inflammatory cells and activated Kupffer cells. In addition, ECM1 heavily decorated apical membranes of BECs forming reactive ductuli in DDC-treated livers, corresponding thus to higher MS intensity of ECM1 during the development of biliary fibrosis. Quantification of IF staining also showed dynamic changes in ECM1 antigen abundance during fibrosis development and resolution confirming thus the MS data (Figure S7E). Induction of ECM1 expression in the inflammatory cells at the sites of primary injury in the CCl_4_ model and in reactive ductuli in the DDC model strongly indicates its involvement in fibrogenesis and healing. Although a previous study had identified ECM1 expression to be hepatocyte-specific,^32^ our data suggest that its role might be more complex than anticipated.

### Correlation of dynamic changes of hepatotoxic proteome and fibrotic deposits identifies clusterin as a novel protein associated with fibrosis resolution

The CCl_4_ model gradually developed typical bridging liver fibrosis^14^ followed by fibrosis resolution as documented by liver injury markers (Figure S1A) and collagen deposits (Figure S1B), as well as by results of our MS profiling (Figure 2). These dynamic changes allowed us to correlate protein abundance profiles (from both Total and E-fraction proteomes) with disease dynamics (captured as fibrotic area) in individual mice (Figure 6). For example, leukocyte-specific integrin β2-interacting protein coronin 1a displayed a significant positive correlation with fibrogenesis reflecting the time-course of the immune response while negatively correlating methionine cycle enzyme adenosylhomocysteinase corresponded to hepatocellular death induced by fibrogenesis (Figure 6B,C).

**FIGURE 6.**
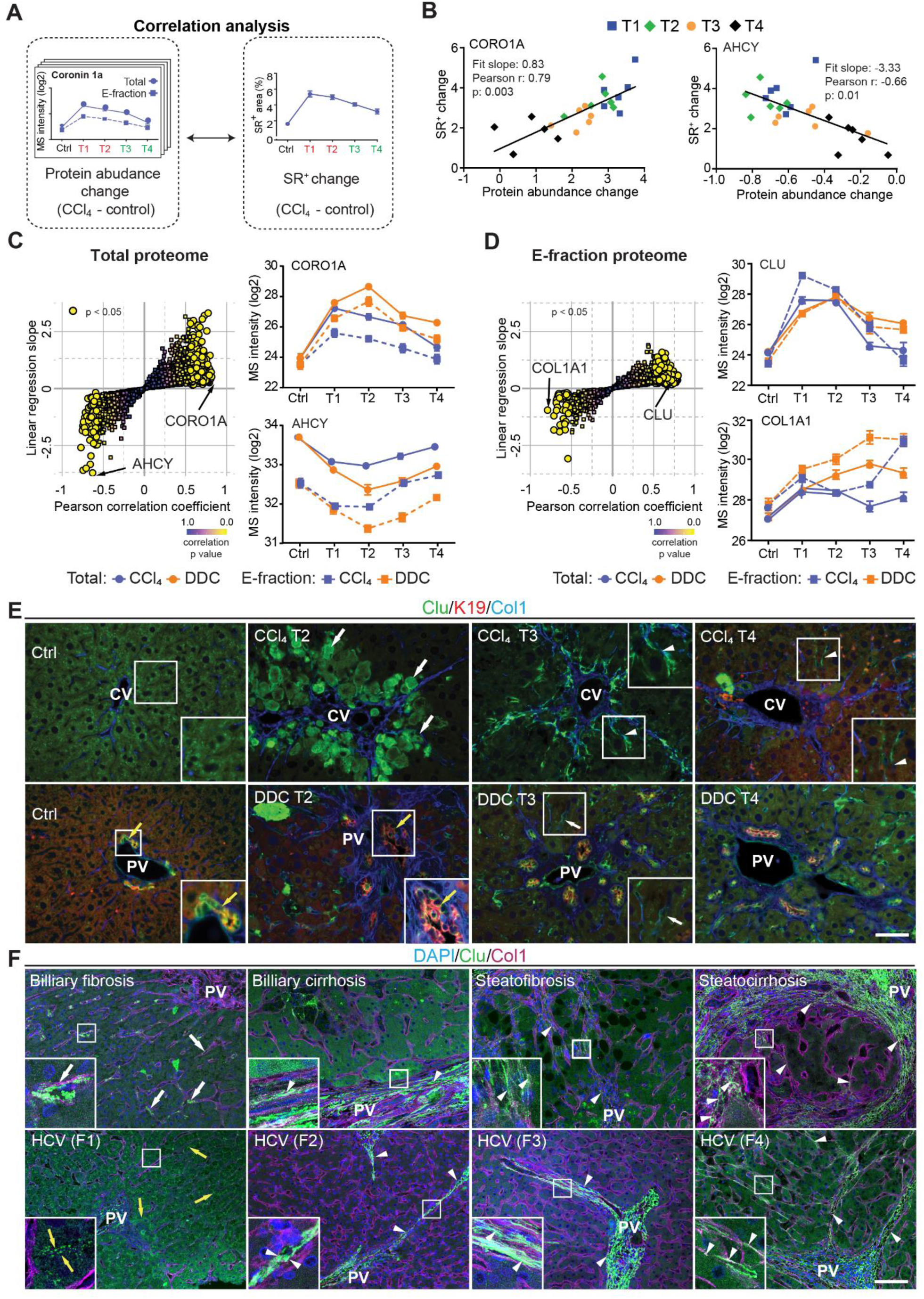
Correlation analysis of protein abundance and changes in fibrotic deposits in the hepatotoxic model associates clusterin with fibrosis resolution. (**A**) Schematic illustrates the correlation of protein abundance changes with changes in sirius red-positive (SR^+^) areas of fibrous ECM deposits in CCl_4_-treated animals at the indicated time points. (**B**) The regulator of the actin cytoskeleton, coronin 1a (CORO1A), serves as an example of a protein with a positive slope of the correlation fit. The methionine cycle enzyme, adenosylhomocysteinase (AHCY), serves as an example of a protein with a negative slope of the correlation fit. (**C,D**) The scatter plots show the linear regression slope and the Pearson correlation coefficient for proteins of CCl_4_ Total (C), and E-fraction (D) proteomes. Statistical significance of the correlation is color-coded as indicated. The line plots show time-dependent change in mass spectrometry intensities of indicated representative proteins with significant correlation in Total (solid line) and E-fraction (broken line) proteomes in CCl_4_ and DDC models; n = 4–6. (**E**) Representative immunofluorescence (IF) images of liver sections from untreated controls (Ctrl), CCl_4_-, and DDC-treated mice at indicated time points of fibrosis development (T2) and resolution (T3 and T4) immunolabeled for clusterin (green), keratin 19 (K19, red), and collagen 1 (Col1, blue). Arrowheads, clusterin staining signal delineating collagen deposits; arrows, clusterin-positive injured hepatocytes; yellow arrows, clusterin-positive biliary epithelial cells. CV, central vein; PV, portal vein. Boxed areas, ×2 images. Scale bar = 50 μm. (**F**) Representative IF images of human liver sections from different stages of chronic liver diseases of various etiologies (biliary-type, steatotic liver disease, and chronic HCV infection) immunolabeled for clusterin (Clu, green) and collagen 1 (Col1, magenta). Nuclei were stained with DAPI (blue). Top row shows increase in clusterin expression along collagen fibrils in biliary-type and metabolic syndrome-related cirrhosis compared to the stage of mild fibrosis. Bottom row documents change in clusterin staining pattern with chronic hepatitis C progression from fibrosis stage F1 to stage F4 (METAVIR grading system: F1, portal fibrosis; F2, periportal fibrosis; F3, bridging septal fibrosis; F4, cirrhosis). Arrowheads, clusterin staining delineating collagen deposits; arrows, clusterin-positive capillarized sinusoids; yellow arrows, clusterin-positive bile canaliculi (stage F1 only). PV, portal vein. Boxed areas, ×4 images. Scale bar = 50 μm.

In the Total CCl_4_ proteome, proteins positively correlating with fibrogenesis included matrisome proteins (e.g., angiotensinogen, fibrinogen α,β, γ-chain, fibrinogen-like protein 1, and fibronectin). The IPA analysis linked these to the inflammation and pathways associated with ECM-cell interactions, such as “Actin cytoskeleton signaling” and “Integrin signaling” (Figure S8A). Proteins with negative correlation were mostly enzymes involved in hepatocyte metabolism, such as alpha-enolase (ENO1), an enzyme shown to participate as plasminogen receptor in ECM degradation.^33^ Analogously to Total proteome, proteins of CCl_4_ E-fraction positively correlating with fibrosis area were linked to fibrogenesis (Figure S8B). Displaying negative correlation was a group of 41 proteins consisting mainly of enzymes associated with amino acid metabolism, necrosis, and apoptosis (Figure S8A,B). In addition, three core matrisome proteins – decorin, MFAP2, and collagen α-1 (I) – also were identified. As shown for collagen α-1 (I), their MS intensity temporal profiles indicated their upregulation in E-fraction while their overall abundance decreased with healing (Figure 6D), likely corresponding to the increased association of these matrisome proteins with degraded insoluble ECM.

Among proteins positively correlating with fibrotic scar development and resolution in both Total and insoluble proteomes was clusterin, a glycoprotein implicated in several diseases (Figure 6D).^34^ As clusterin imparts extracellular chaperone function not previously linked to fibrosis, we next investigated clusterin’s localization by IF microscopy (Figure 6E). In untreated livers, clusterin was detected in BECs and partially in hepatocytes. Upon CCl_4_ intoxication, IF staining revealed its prominent expression by damaged hepatocytes. Strikingly, clusterin increasingly re-localized along the collagen deposits over the healing period. In DDC-treated livers, clusterin was found mostly in BECs forming reactive ductuli and hepatocytes, with minimal changes upon healing (Figure 6E). Moreover, extracellular clusterin deposition in close proximity of collagen fibers was further demonstrated in fibrotic and cirrhotic human livers of various etiologies at different stages of fibrosis (Figures 6F and S9). The gradual increase in clusterin expression with the progression of fibrosis against the background of chronic hepatitis C (METAVIR score F1-F4) further emphasized its role in the development of liver fibrotic diseases.

### Interface hepatocyte elasticity responds dynamically to the pericentral injury in the course of fibrosis development

Identification of “Rho-A”, “Rho family GTPases”, “Actin cytoskeleton”, and “Integrin/ILK signaling” among top canonical pathways elicited by proteins of Total CCl_4_ proteome as well as proteins of matrisome-enriched clusters (Figure S3A,C) suggested that the hepatotoxic injury leads to increased cytoskeletal tension in injured hepatocytes. This was further supported by significant enrichment in intermediate filament proteins (Figure 2E) and in myocardin-related transcription factor (MRTF) targets (e.g., ITGA1, THBS1, ACTR2, and MSN; Figure 7A), key regulators involved in cell and tissue mechanics, cytoskeletal dynamics, and mechanosensing.^35^

**FIGURE 7.**
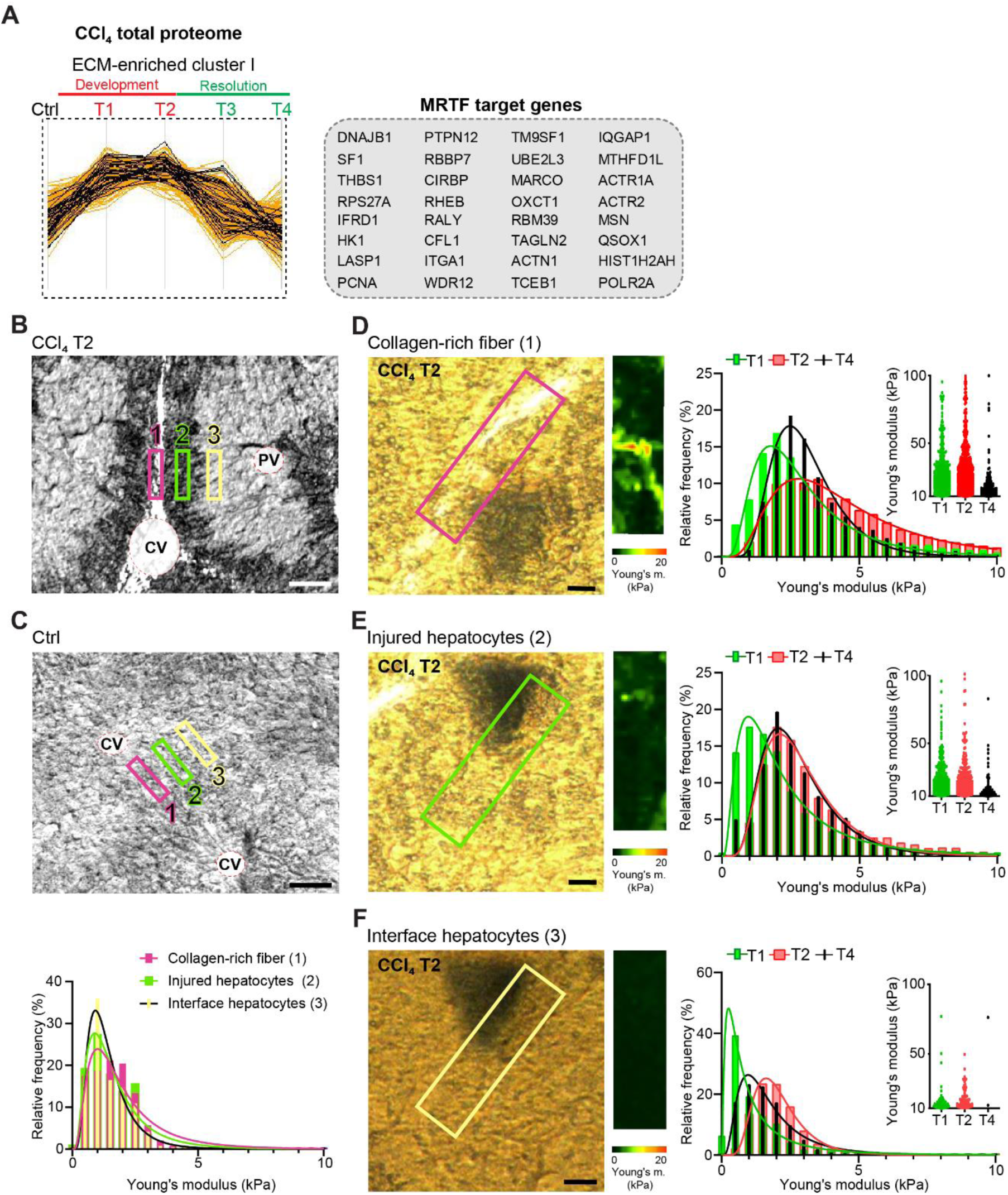
Atomic force microscopy (AFM) stiffness mapping reveals changes in the local mechanics of liver tissue upon hepatotoxic injury. (**A**) The line plot shows the dynamics of 32 myocardin-related transcription factor (MRTF) targets (highlighted in black) significantly enriched in ECM-enriched cluster I of CCl_4_ Total proteome (Figure 2C) identified by Fisher’s exact test (*p* = 0.01; Benjamini-Hochberg FDR = 0.03). (**B,C**) Representative polarized microscopy images of CCl_4_-treated (T2, B) and untreated control (Ctrl, C) liver sections with indicated regions (1–3) selected for AFM measurements. Note white areas corresponding to collagen fibers visualized by polarized light. Pink rectangle, region 1 (collagen-rich fiber); green rectangle, region 2 (injured hepatocytes); yellow rectangle, region 3 (interface hepatocytes). CV, central vein; PV, portal vein. Scale bar = 100 μm. Histogram shows Young’s moduli for the measured regions in untreated control livers; n = 7 regions in three mice. (**D–F**) Representative polarized microscopy images of CCl_4_-treated (T2) liver sections with rectangle indicating the regions of AFM measurements (30 × 100 μm^2^) and corresponding pseudocolor Young’s modulus maps determined by AFM. Scale bar = 25 μm. Histograms show Young’s moduli for indicated regions of collagen-rich fibers (D), injured hepatocytes (E), and interface hepatocytes (F) at indicated time points of fibrosis development (T1 and T2) and spontaneous resolution (T4). Inset scatter plots show Young’s modulus values above 10 kPa for each time point; n = 7 regions in three mice.

As variations in ECM composition and content also substantially affect the biomechanics of liver tissue, we decided to examine correlation between our proteomic results and mapping of dynamic changes in the biomechanical properties of CCl_4_-treated livers. We used atomic force microscopy (AFM) combined with polarized microscopy^17^ to precisely locate and probe the following compartments: 1) regions of collagen-rich scar tissue in close vicinity of the central vein, 2) injured hepatocyte regions next to the collagen scar, and 3) regions of hepatocytes on the interface between injured and not visibly damaged hepatocytes (so-called interface hepatocytes^36^) (Figure 7B). In parallel, we also measured stiffness of corresponding regions in nontreated control liver sections (Figure 7C).

Our AFM analysis revealed no substantial topological variations across defined compartments in control livers, with median Young’s moduli ranging between 1.2 and 1.6 kPa (Figure 7C, Table S1). Fibrogenic response triggered progressive tissue stiffening apparent in all analyzed compartments (Figure 7D–F, Table S1), with collagen-rich regions the stiffest at maximal fibrosis (∼4.4 kPa at T2; Figure 7D). Unexpectedly, interface regions exhibited initial softening (T1; median ∼0.8 kPa vs. 1.2 kPa in control livers), which was followed by increase in stiffness up to ∼2.0 kPa (T2). With healing (T4), all compartments demonstrated clear decrease in stiffness (Figure 7D–F), although this was only marginal in the regions corresponding to injured hepatocytes. Significant softening of collagen-rich scar tissue during healing indicated partial resolution of scar-associated ECM while it associated with the incorporation of several core matrisome proteins (e.g., COL1A1 and FBN1) into the scar tissue (Figures 1F and 6D). Although the heterogeneity of AFM-based measurements constituted a limitation on our correlation analysis relative to proteomic data (Figure S10), this analysis provides a coherent framework for better understanding the dynamics of proteomic landscapes in the context of fibrosis-associated changes in the local mechanics of liver tissue.

## DISCUSSION

In this study, we comprehensively characterized and compared dynamic proteomic landscapes of liver fibrosis development and repair in two mouse models based either on repetitive CCl_4_ intoxication^14^ or DDC feeding.^16^ These models are widely used to reliably mimic either hepatotoxic (CCl_4_) or cholestatic (DDC) liver injuries leading to fibrosis in humans.^14^ Previous attempts to grasp the complexity of etiology-specific fibrotic proteomes with a focus on diseased ECM^6, 7^ were limited due to origins of the analyzed material (mostly decellularized samples) and yielded only fragmented insight into the time course of the disease. To fill this research gap, we employed detergent-based tissue extraction and analyzed both Total and insoluble fraction proteomes at multiple time points of disease progression and spontaneous healing. This approach allowed us to 1) compare time-resolved, compartment-specific proteomes of CCl_4_ and DDC models, 2) define etiology-specific elements of fibrogenesis, and 3) detect even low-abundance matrisome and matrisome-interacting proteins in the ECM-enriched insoluble fraction samples. Further, our detailed AFM-based profiling of CCl_4_-treated livers enabled us to relate our proteomic data to disease-associated changes in the local mechanics of liver tissue.

We and others have demonstrated that the nature of liver injury determines the set of components assembled in diseased ECM,^7^ which thus reflects unique features of the injury. For instance, the initial decline in abundance of hepatocyte- and activated BEC-expressed COL18A1^37^ and its consecutive increase thereafter in the hepatotoxic CCl_4_ model corresponds to hepatocyte death followed by expansion of both COL18A1-expressing cell types.^38^ By contrast, extensive COL18A1 expression in DDC-induced cholestasis was already apparent in the early fibrotic deposits due to a massive ductular reaction. A prominent group of BM-associated proteins uniquely identified in the cholestatic model can be attributed to activated BECs in proliferating bile ducts. Some of them (e.g., COL18A1, COL4A2, COL6A2, and HSPG2) are among the 14-gene signature potently predicting human patient cirrhosis progression and survival.^39^ Other DDC matrisome constituents (e.g., COL3A1, elastin, and LAMC1) have been recently linked to the deterioration of hepatocyte functions in connection with ECM stiffening.^40^ Notably, elastin is present in liver biopsies from patients with advanced fibrosis and adverse clinical outcome^41^ and is associated with the irreversibility of liver fibrosis. Further, the increased incorporation of crosslinking protein LOXL1 and TGFβ1-related protein LTBP4 into DDC-specific ECM underscores its resemblance to human cirrhotic liver ECM.^42^ Hence, our findings indicate that cholestasis-driven ECM deposits contain numerous proteins detected in more advanced stages of liver disease favoring hepatocarcinogenesis with a compromised ability to heal. Moreover, identified cholestasis-induced unique signaling pathways were analogous to those recently identified in a subtype of mouse cancer models mimicking human cholangiocarcinoma-like hepatocellular cancer.^43^

Our observations are concordant with recently published studies (reviewed in^2^) revealing that in the fibrotic liver resident and nonresident cell types wire into dynamic intercellular hubs with shifting cell populations, reflecting and shaping the disease course in an etiology-specific manner. Many of the feedback mechanisms between the cells and ECM regulating fibrosis are mediated by members of the integrin family. In agreement with previous studies,^27^ we observed in both models significant upregulation of all detected integrin subunits during fibrosis progression. Strong induction of integrin αM expression in cholestatic but not hepatotoxic injury documents the role of leukocyte-specific integrin αMβ2 in the modulation of biliary fibrosis,^44^ suggesting that the integrin αM might thus serve as a possible selective target for treatment of cholestatic liver disease. Further, a prominent transition of TGFβ1-activating integrin αv toward insoluble ECM exclusively upon hepatotoxic injury was accompanied by its localization to injured pericentral hepatocytes near collagen-rich scars and αSMA-positive HSCs. This finding, together with a measured increase in the stiffness of injured hepatocytes, implies an active role of integrin αv in the targeted activation of TGFβ1 in the local microenvironment during centrilobular fibrosis. Indeed, integrin αv located on hepatocytes in the vicinity of biliary fibrotic septa has been suggested to indicate hepatocyte biliary transformation.^45^ Here, we propose that compartment-specific induction of αv expression on the surface of scar-associated hepatocytes could be a general mechanism to promote fibrosis progression.

In contrast to the cholestatic DDC model, hepatotoxic injury in the CCl_4_ model showed a substantial ability to heal 10 days after challenge withdrawal. Such healing capacity allowed us to correlate the dynamics in CCl_4_ proteomes with the changes in fibrotic deposits not only during fibrosis progression but also in resolution. Among proteins correlating with scar tissue formation in both Total and insoluble proteomes, we identified a small molecular chaperone clusterin. Clusterin is believed to be associated with elastin fibers in cholestasis,^46^ and recently it was linked to the attenuation of mouse hepatic fibrosis.^47^ Here, we localized clusterin exclusively to injured, pathologically stiffer hepatocytes. Interestingly, the clusterin promoter comprises binding motifs for mechanosensitive transcription factors such as Fos and AP-1/Jun.^48, 49^ This suggests that local clusterin induction can reflect the dynamic changes of microenvironmental mechanical cues. Most intriguingly, we also found clusterin to associate with collagen-rich ECM deposits in the course of mouse fibrosis regression and with pathological human matrix of various etiologies. As clusterin overexpression associates with increased activity of ECM-degrading matrix metalloproteinases,^50^ we hypothesize that clusterin accumulation facilitates the remodeling and resolution of scar tissue. Although the specific molecular function of clusterin in fibrotic tissue repair processes remains to be determined, our results strongly suggest clusterin to be an attractive antifibrotic target.

Liver cell and tissue mechanics play a pivotal role in the processes that initiate and resolve fibrotic injury.^3^ Initial disruption of mechanical homeostasis prompts cytoskeletal remodeling that alters cell-generated forces and cellular biomechanics. Here, such a shift to a higher stiffness regime is illustrated by prominent changes in cytoskeleton-related signatures (actin and intermediate filaments) and signatures of mechanosensitive transcription regulators (e.g., MRTF) accompanied by a significant stiffening of parenchymal compartments devoid of apparent collagen deposits. This indicates substantial cell-driven changes in the biomechanical properties of tissue microenvironment. An unexpected decrease in the stiffness of interface hepatocytes with the initial fibrotic changes revealed by our AFM analysis suggests that the initial softening of interface hepatocytes upon injury counterbalances the stiffening of the injured hepatocytes and/or fibrous scar tissue regions. As the interface hepatocytes have been shown to undergo a phenotypic shift in response to the injury,^36^ it will now be interesting to determine whether altered mechanical properties serve as the cue leading to the genetic fetal reprogramming or if this initial softening is due to the expression of fetal markers. Together, our AFM and proteomic data underscore the role of local tissue stiffening in fibrotic response and postulate involvement of tissue softening during resolution not only in the collagen scar tissue but also in regenerating parenchyma.

## Author Contributions

Conceptualization: M.G. and M.J. Acquisition of data: M.J., K.H., S.O., K.K., and L.S. Analysis and interpretation of data: M.J., K.H., P.C., S.O., and M.G. Drafting of the manuscript: M.J. and M.G. Critical revision of the manuscript for important intellectual content: all authors. Funding: M. G. and M.J. Technical and material support: H.B.S., E.S., and C.H.M.

## List of Abbreviations

AFM: atomic force microscopy
BECs: biliary epithelial cells
B.H. FDR: Benjamini-Hochberg false discovery rate
CCl_4_: carbon tetrachloride
DDC: 3,5-diethoxycarbonyl-1,4-dihydrocollidine
ECM: extracellular matrix
ECM1: extracellular matrix protein-1
GO term: gene ontology
HSC: hepatic stellate cell
IPA: Ingenuity Pathway Analysis
MRTF: myocardin-related transcription factor
MS: mass spectrometry
PCA: principal component analysis
PF: portal fibroblasts

## ACKNOWLEDGEMENTS

We would like to thank S.M. Meier-Menches (University of Vienna, Austria) and J. Masek (Charles University, Prague, Czech Republic) for critical reading of the manuscript; and D. Hadraba (Institute of Physiology, Prague, Czech Republic) for sharing his expertise. We further acknowledge the Light Microscopy Core Facility, IMG CAS, Prague, Czech Republic for support with the microscopy imaging presented herein. We would also like to thank V. Getmanchuk - Zaporoshchenko for detailed proofreading of the final manuscript.

This work was supported by the Grant Agency of the Czech Republic (18-02699S and 21-21736S); the Grant Agency of the Ministry of Health of the Czech Republic (NU21-04-00100); the Institutional Research Project of the Czech Academy of Sciences (RVO 68378050); the National Institute for Cancer Research (Programme EXCELES, LX22NPO5102) - Funded by the European Union - Next Generation EU; the Grant Agency of Charles University (273723), and MEYS CR projects (LM2023050, LM2018126, LQ1604 NPU II, LO1419, and LM2015040). CIISB, Instruct-CZ Centre of Instruct-ERIC EU consortium, funded by MEYS CR infrastructure project LM2018127 and European Regional Development Fund-Project “UP CIISB” (No. CZ.02.1.01/0.0/0.0/18_046/0015974) financially supported the measurements at the CF Nanobiotechnology, CEITEC MU (AFM measurements). The funding sources were not involved in the study design, data collection and analysis, decision to publish, or preparation of the article.

Presentation: none.

## DATA AVAILABILITY STATEMENT

The data that support the findings of this study are available from the corresponding author upon reasonable request.

