## Supplemental Material for "Dynamics of compartment-specific proteomic landscapes of hepatotoxic and cholestatic models of liver fibrosis"

**This file contains:**

**1. Supporting MATERIALS AND METHODS**

**2. Supporting RESULTS**

**3. Supporting REFERENCES**

**4. Supporting FIGURES**

**5. Supporting TABLES**

### 1. Supporting MATERIALS AND METHODS

#### ETHICAL STATEMENT

##### Human samples

The use of completely anonymized archived liver tissue samples for research purposes has been approved by the Ethical Committee of the Institute of Experimental and Clinical Medicine and Thomayer University Hospital, Prague, Czech Republic. Written informed consent was obtained from all patients enrolled in the study. All research was conducted in accordance with both the Declarations of Helsinki and Istanbul.

##### Animals

Male C57BL/6J mice (8-10 weeks old) were housed under standard pathogen-free conditions with free access to regular chow and drinking water and a 12-hour-dark/12-hour-light cycle. All animal experiments were performed in accordance with an animal protocol approved by the Animal Care Committee of The Institute of Molecular Genetics and according to EU Directive 2010/63/EU for animal experiments.

#### LIVER INJURY MODELS

To induce fibrogenesis, mice were either injected intraperitoneally with 1 $\mu$l/g carbon tetrachloride ($CCl_4$; diluted 1:3 with olive oil) twice a week for 3 or 6 weeks or fed a diet supplemented with 0.1% 3,5-diethoxycarbonyl-1,4-dihydrocollidine (DDC; Sniff, Soest, Germany) for 2 or 4 weeks. To study fibrosis resolution, mice treated with $CCl_4$ for 6 weeks or fed DDC for 4 weeks were removed from treatment and allowed to recover for 5 or 10 days. At the end of treatment, mice were anesthetized, blood was collected into heparin-containing tubes (KabeLabortechnik, Numbrecht, Germany) to obtain plasma, and mice were sacrificed by cervical dislocation to collect liver samples. Plasma levels of aspartate transaminase (AST), alanine transaminase (ALT), alkaline phosphatase (ALP), and total bilirubin, were measured using commercial kits (Roche Diagnostics, Prague, Czech Republic).

A 1 mm thick middle section of left lateral liver lobe was removed and two ~1 $mm^3$ pieces were cut, flash frozen in liquid nitrogen, and stored at -80°C for further processing for mass spectrometry (MS). The rest of the lobe was divided into two parts; one was fixed in 4% buffered formalin (pH 7.4) followed by paraffin embedding for histology and immunostaining and the other was embedded in OCT Tissue-Tek (Sakura Finetek, USA) for further cryosectioning for AFM measurements.

**HISTOLOGY AND IMMUNOFLUORESCENCE**

Formalin-fixed, paraffin-embedded liver sections (4 µm; both mouse and human) were stained with hematoxylin and eosin (H&E), and sirius red (SR). For immunofluorescence, these sections were de-paraffinized and subjected to heat-induced antigen retrieval in Tris-EDTA (pH 9) or citrate (pH 6) buffer and further permeabilized with 0.1 M glycine, 0.1% Triton X-100 (15 minutes). Next, sections were incubated with primary antibodies overnight at 4°C, followed by incubation with secondary antibodies for 120 minutes at 22°C. The following primary antibodies were used: rat mAb to and K19 (Troma III; Developmental Studies Hybridoma Bank, University of Iowa, Iowa City, USA); mouse mAbs to αSMA (DAKO, Glostrup, Denmark) and ECM1 (Santa Cruz Biotechnology, Dallas, USA), goat polyclonal antibodies to clusterin (R&D Systems, Minneapolis, USA), rabbit polyclonal antibodies to fibronectin, collagen I, collagen IV, and monoclonal to integrin αV (all from Abcam, Cambridge, UK). As secondary antibodies we used donkey anti-mouse IgG Alexa Fluor (AF) 488 and 647, goat anti-mouse IgG and goat anti-rabbit IgG horseradish peroxidase (HRP)-conjugated, donkey anti-rabbit IgG AF488, donkey anti-rat IgG AF594 (all from Jackson ImmunoResearch, Baltimore, USA). ECM1 and integrin αV were visualized using Alexa Fluor™ 488 Tyramide Reagent (ThermoFisher Scientific, Prague, Czech Republic) according to the manufacturer's instructions. Stained sections were further mounted with ProLongGold Antifade mounting media (ThermoFisher Scientific) and images were acquired with inverted fluorescence microscope DM6000 (Leica Microsystems, Pragolab, Prague, Czech Republic) equipped with HC PL Apo 10×/0.40 NA, PH1 HC PLAN Apo 20×/0.70 NA, and PH2 HCX PL Apo 40×/0.75 NA, objective lenses.

**MASS SPECTROMETRY SAMPLE PREPARATION**

A sample of ca 1 $mm^3$ excised from the middle section of left lateral liver lobe was cryo-homogenized using TissueLyser II (QIAGEN, Germantown, MD, USA). The cryo-homogenized liver powder was then resuspended in 200 µl of 50 mM Tris-HCl pH 7.5, 5% glycerol, 500 mM NaCl, 1% IGEPAL CA-630, 2% sodium deoxycholate, 1% SDS, 1× Proteinase Inhibitor Cocktail. The suspension was sonicated after 20 minutes incubation at 22°C. A 50 µl aliquot of the resuspended sample was further precipitated with ice-cold acetone and labeled „Total". The remaining solution was centrifuged at 16,000 × g for 20 minutes to separate the supernatant containing soluble (S)-fraction from the pellet, which itself contained insoluble ECM-enriched (E)-fraction. Supernatants were further precipitated with ice-cold acetone and stored together with pellets at -80°C for further processing preceding MS analysis.

**PROTEIN DIGESTION**

Protein pellets were homogenized and lysed in 100 mM TEAB (Triethylammonium bicarbonate) containing 2% sodium deoxycholate, 40 mM chloroacetamide, 10mM TCEP (Tris(2-carboxyethyl)phosphine) at 95°C for 10 minutes followed by sonication (Bandelin Sonopuls Mini 20 homogenizer). Protein concentration was determined using a BCA protein assay kit (ThermoFisher Scientific) and 30 µg of protein per sample was used for MS sample preparation as described before[1]. Briefly, 5 µl of SP3 beads were added to 30 µg of protein sample and brought to 50 µl with 100 mM TEAB. Protein binding was induced by the addition of ethanol to 60% (v/v) final concentration. Samples were mixed and incubated for 5 minutes at 22°C. After binding, tubes were placed into the magnetic rack and the unbound supernatant was discarded. Beads were subsequently washed two times with 180 µl of 80% ethanol. Samples were further digested with trypsin (trypsin/protein ratio 1/30, reconstituted in 100 mM TEAB) at 37°C overnight followed by acidification with trifluoroacetic acid to 1% final concentration. Peptides were desalted using in-house-made stage tips packed with 3M™ Empore™ C8 extraction disks (Thermo Fisher Scientific) as described before[2].

**nLC-MS/MS ANALYSIS**

Nano Reversed phase columns (EASY-Spray column, 50 cm × 75 µm ID, PepMap C18, 2 µm particles, 100 Å pore size) were used for LC/MS analysis using 0.1% formic acid in water as mobile phase buffer A and 0.1% formic acid in acetonitrile as mobile phase B. Samples were loaded onto the trap column (C18 PepMap100, 5 µm particle size, 300 µm × 5 mm; Thermo Fisher Scientific) for 4 minutes at 18 µl/minute in loading buffer (2% acetonitrile, 0.1% trifluoroacetic acid). Peptides were eluted with mobile phase B gradient from 4% to 35% in 120 minutes. Eluting peptide cations were converted to gas-phase ions by electrospray ionization and analyzed on a Thermo Orbitrap Fusion (Q-OT-qIT, Thermo Scientific). Survey scans of peptide precursors from 350 to 1400 m/z were performed in orbitrap at 120 K resolution (at 200 m/z) with a $5 \times 10^5$ ion count target. Tandem MS was performed by isolation at 1,5 Th with the quadrupole, HCD fragmentation with a normalized collision energy of 30, and rapid scan MS analysis in the ion trap. The MS2 ion count target was set to $10^4$ with the max injection time of 35 milliseconds. Only those precursors with charge state 2-6 were sampled for MS2. The dynamic exclusion duration was set to 45 seconds with a 10 ppm tolerance around the selected precursor and its isotopes. Monoisotopic precursor selection was turned on. The instrument was run in top speed mode with 2 seconds cycles[3].

### MS INTENSITY DATA QUANTIFICATION

Peptides were identified and quantified by MaxQuant label-free quantification software (1.6.7 version[4]) using *Mus musculus* UniProt protein database (UniProtKB version July 2020, containing 25,805 entries) and the instrument type set to Orbitrap. Trypsin was set as the proteolytic enzyme and two missed cleavages were allowed. Carbamidomethylation of cysteine was selected as a fixed modification and N-terminal protein acetylation and methionine oxidation as variable modifications. Reverse sequences were selected for the target-decoy database strategy[5]. The false discovery rate (FDR) was set to 1% for both proteins and peptides a minimum peptide length specified to seven amino acids. The "match between runs" feature of MaxQuant was used to transfer identifications to other LC-MS/MS runs based on their masses and retention time and this was also used in quantification experiments. Quantifications were performed with the label-free algorithm in MaxQuant[6]. The Fast LFQ option was switched off. Each sample of total protein extract was treated as a separate 'experiment' for quantification. Corresponding E- and S-fractions together were treated as another separate 'experiment'. Unique+razor values were used for LFQ quantification. Protein groups identified through target-decoy database strategy ('Reverse' column in the MaxQuantproteinGroups file) and proteins not identified by MS/MS spectrum ('Only identified by site' column in the MaxQuantproteinGroups file) were removed and not included in any subsequent analysis.

The overall protein intensity for total extract samples was estimated as a sum of MaxQuant label-free quantification values (LFQ) for unique and razor-detected peptides. For each S-fraction sample and corresponding E-fraction sample, the proportional coefficients for LFQ quantification values of unique and razor-detected peptides of each 'experiment' were computed according to Cox et al.[6] using the script for R software (version 4.0.5 for Windows) and the values in the MaxQuant evidence file. After the estimation of the proportional coefficients for each fraction, the overall protein MS intensity for each fraction was estimated as a sum of normalized LFQ values for unique and razor-detected peptides.

### BIOINFORMATIC ANALYSIS AND STATISTICS

Normalized[7,8] log 2-transformed MS intensity data quantified as described above were uploaded for bioinformatic analysis into the bioinformatics platform Perseus (1.6.10.43 version[9]). The used annotations were: gene ontology (GO), biological process (BP), molecular function (MF), cellular component (CC), KEGG pathway databases, matrisome database[10, 11], previously published cell type-specific protein and gene expression signatures of liver cells[12]and leukocyte populations[13-15], annotations „location“ and „function“ from Ingenuity Pathway Analysis[16] annotation (IPA, QIAGEN Inc., https://www.qiagenbioinformatics.com/products/ingenuitypathway) and myocardin-related transcription factor (MRTF) target genes[17]. All data analyzed were first log 2 transformed. The PCA analysis was performed in Perseus software with the built-in tool and all data imputed by random selection from a normal distribution generated at 1.8 standard deviations subtracted from the mean of the total intensity distribution and a width of 0.3 standard deviations. For Total proteome sample analysis, data from Total samples were first filtered for proteins expressed in at least one group (control, and times T1, T2, T3, or T4) with at least 5 identified values in both models ($CCl_4$ and DDC) together. For subsequent analyses we also similarly filtered Total proteome samples in each model separately. Missing values among identified proteins in analyzed data sets were imputed into Perseus as described above. Proteins significantly differentially expressed in the Total proteome of each fibrosis model separately (analyses in Figure 2 and 3) were defined as proteins with a minimum 1.5 fold significant change in expression in the highest fibrosis time point (T2) over the average expression in controls (t-test at a Benjamini-Hochberg (B. H.) false discovery rate (FDR) < 0.05). Unsupervised hierarchical cluster analyses of averages of z-scored MS intensities of identified proteins were performed with Euclidean distance and complete linkage in the Perseus software with built-in cluster enrichment analysis. To extract individual cell-type dynamics from the proteomics data we used previously published cell-type-specific signatures [12-15]. For each of this way identified proteins, average z-score MS intensity in each time point was calculated and the signature proteins of each cell type were together plotted in each time point as box plots, with 25th and 75th percentile indicated as explained below.

All graphs and statistical tests indicated in graphs were performed using GraphPad Prism version 9.3.1 (GraphPad Software, Boston, Massachusetts USA, www.graphpad.com). All results are presented as mean ± SEM. In the boxplots, the box represents the 25th and 75th percentile with the median indicated; whiskers reach the last data point; dots indicate means of independent experiments. Normally distributed parametrical data were analyzed by two-tailed unpaired student's t test. A comparison between multiple groups was performed using one-way ANOVA with Tukey's multiple comparison test. Data distribution was assumed to be normal, but this was not formally tested. Statistical tests used are specified in the figure legends. Statistical significance was determined at the level of *, $p < 0.05$; **, $p < 0.01$; ***, $p < 0.001$.

**E-FRACTION ANALYSIS**

Proteins identified in E-fraction samples were first filtered and imputed as described above for each model separately. All proteins detected in S-fraction were also imputed in Perseus as described above and median S-fraction MS intensity value for each protein was calculated. To identify proteins with change in solubility, which would be specifically enriched in E-fraction, we calculated a ratio of the MS intensity of each protein replicate in E-fraction to the median of MS intensity of this protein in S-fraction and selected those with a minimum threefold higher expression in E- than S-fraction in at least one time point (control, T1-T4). From these, proteins with a significant change in time identified by ANOVA with B. H. FDR < 0.05 represented insoluble fraction proteins, i.e. E-fraction proteome. Unsupervised hierarchical clustering of MS intensity ratios of the abundance of these proteins in E- to S-fraction was performed in Perseus as described above to identify changes in protein solubility over time with disease progression and recovery.

**INGENUITY PATHWAY ANALYSIS**

We used the Ingenuity Pathway Analysis[16] to identify predicted upstream transcriptional regulators and growth factors and downstream canonical signaling pathways elicited by groups of proteins identified in individual analyses. Transcriptional regulators and growth factors were identified in upstream regulator analysis with protein overlap *p*-value > 3 (log 10) and activation z-score > |2.5|. Canonical pathways were considered only with overlap *p*-value > 3 (log 10) and activation z-score > |2|.

**CORRELATION ANALYSIS**

The MS protein temporal abundance change profile of each protein was correlated with the fibrosis area change for proteins of Total proteome and E-fraction proteome in each model. To do this, we calculated the mean value of log 2 transformed and imputed MS intensities for each protein in controls and subtracted it from the log 2 transformed MS intensity for each protein at each timepoint in individual mice. This way obtained temporal abundance profile ratio of each protein was then correlated to the temporal profile of fibrosis area (calculated as a percentage of Sirius red positive areas in each mouse at each timepoint after subtracting the background calculated as the average Sirius red positive area in control livers) using Pearson correlation coefficient separately for each model. We then plotted the correlation linear regression slope against the Pearson correlation coefficient in each of the proteomes for each of the identified proteins (Figure 6B-D). The significance of dependency (slope's *p*-value) for each protein was corrected by B. H. multiple comparisons test (*p* = 0.05). Analysis of DDC-induced differentially expressed proteins did not result in any proteins significantly correlating with collagen deposition in either of the proteomes (Figure S11). This is most likely a consequence of considerable variability in the scar tissue deposition in this model at each time point.

### ATOMIC FORCE MICROSCOPY

Atomic force microscopy (AFM) was performed on 30 μm thick liver sections shortly fixed with 4% PFA in PBS, as previously described[18]. Briefly, glass slides containing fixed liver sections covered with PBS were placed onto a combined AFM/inverted optical microscope (JPK NanoWizard 3XP; Bruker Nano Nano Surfaces Division, Santa Barbara, CA, USA/Olympus IX81; Olympus C&S, Prague, Czech Republic) equipped with a custom made dual-polarization motorized filter allowing precise localization areas of interest: collagen-rich fibrotic areas, areas of injured and interface hepatocytes[19]. To characterize the mechanical properties of livers, we used an SD-qp-BioT-TL-10 cantilever (S/N-73750F05, Nanosensors, NanoAndMore GmbH, Wetzlar, Germany) with a nominal sprint constant of 0.09 N/m modified with a 5.7 μm melamine polymer bead (Microparticles GmbH, Germany). Before each experiment, the cantilever spring constant was calibrated using the thermal noise method while immersed in PBS. Using the force mapping method, we measured 30 ×100 $\mu m^2$ (10 × 36 pixels) defined areas precisely located by polarized microscopy. We analyzed seven areas (measured from 2 sections per mouse) obtained from 3 different mice for each time point (control, T1, T2, and T4). Data were further processed with open-source software "AtomicJ"[20] using a Poisson ratio of 0.45[21]. All data were further analyzed in GraphPad Prism software and are presented as frequency distribution histograms. Statistical analyses were performed on frequency distribution histograms using the Kruskal-Wallis test followed by Dunn's multiple comparisons post-testing. To correlate the AFM data with protein MS abundance change during fibrosis development and resolution we averaged median Young's moduli for each analyzed area in each measured mouse. After subtraction of mean control Young's moduli, the slope and P-value for the dependency of the stiffness of the area on the protein MS intensities were determined, and the P-value was corrected by B. H. multiple comparisons test ($p < 0.05$).

### 2. Supporting RESULTS

#### Mass spectrometry data characterization

MS analysis quantified 4,737 proteins in all liver samples (Figure 1B). The numbers of quantified proteins were comparable across time and treatments. The proteins were equally represented in each of the cohorts. No bias was shown against any of the analyzed cohorts or time points (Figure 1B). Likewise, the numbers of quantified proteins across fractions showed only minor variability (Figure 1C). We identified 184 matrisome constituents[10, 11] among the quantified proteins. Their median sequence coverage in controls, $CCl_4$, and DDC-samples only slightly differed from sequence coverage of all identified proteins, showing no strong bias against matrisome-annotated proteins (Figure 1C). Principal component analysis (PCA) demonstrated the separation of $CCl_4$- and DDC-derived samples (Figure 1D), as well as the separation of Total and E-fraction samples in both $CCl_4$ and DDC models in the first two components (Figure 1E). Samples also separated in time, which demonstrated fibrosis development and its partial resolution for Total and E-fraction samples in both models (Figures 1E).

#### Qualitative changes in total proteomes induced by $CCl_4$ and DDC models

Liver injury elicits not only changes in protein abundance but also results in qualitative changes in proteome. To analyze proteins that are induced during fibrosis development in Total proteome we identified proteins detected in T2 and T1 but not in control samples in each model. We identified 65 newly induced proteins in $CCl_4$- and 118 in DDC-treated livers. These were mainly matrisome proteins and proteins of the immune defense response (Figure S3D). Notably, we found lysyl hydroxylase (Plod3), a matrisome protein highly expressed in activated HSCs[22], among newly expressed $CCl_4$-specific proteins. In contrast, DDC-specific matrisomal proteins included elastin, a proposed portal fibroblast marker[23] (Figure S3D). This confirmed the specificity of identified $CCl_4$ and DDC signatures and suggested the potential of our approach in describing the different origin of collagen-producing myofibroblasts in analyzed models.

**4. Supporting FIGURES**

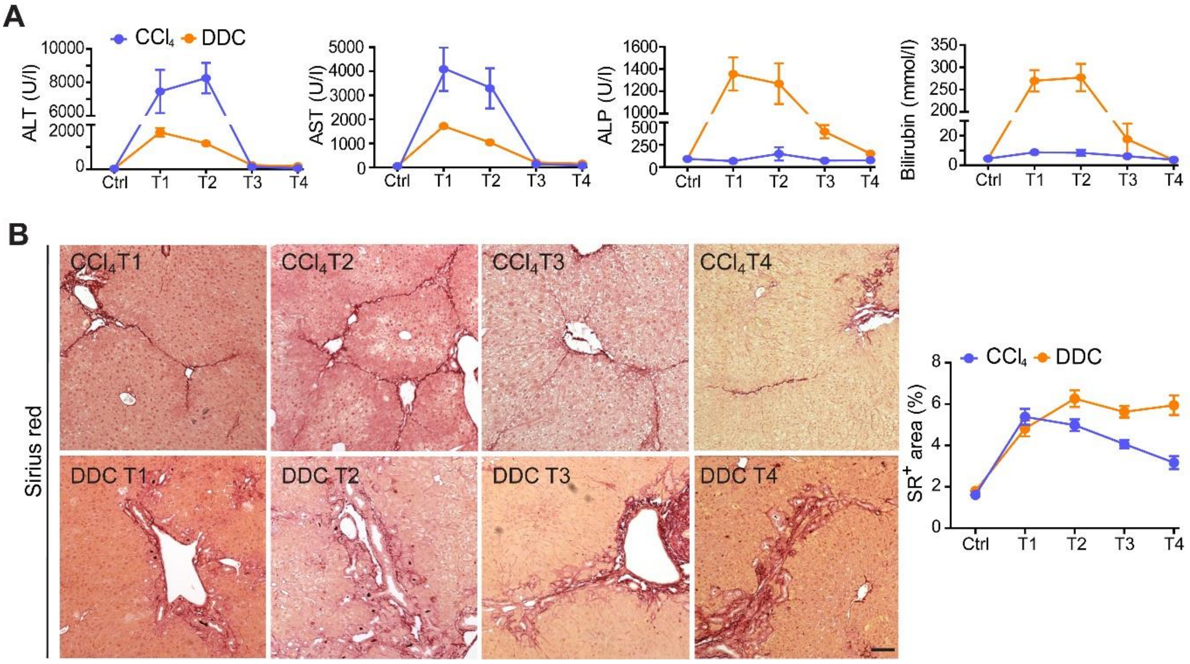


**FIGURE S1** Liver damage markers and collagen deposits in $CCl_4$ and DDC models. (**A**) Serum levels of alanine aspartate transaminase (ALT), aspartate transaminase (AST), alkaline phosphatase (ALP), and bilirubin determined from $CCl_4$- and DDC-treated mice at indicated time points. (**B**) Representative images of SR-stained liver sections from mice treated with $CCl_4$ or DDC document time-course in fibrosis development and partial resolution. Scale bar = 50 μm. Line plot shows the quantification of collagen deposits in each time point shown as a percentage of the $SR^+$ area per liver section; n = 6.

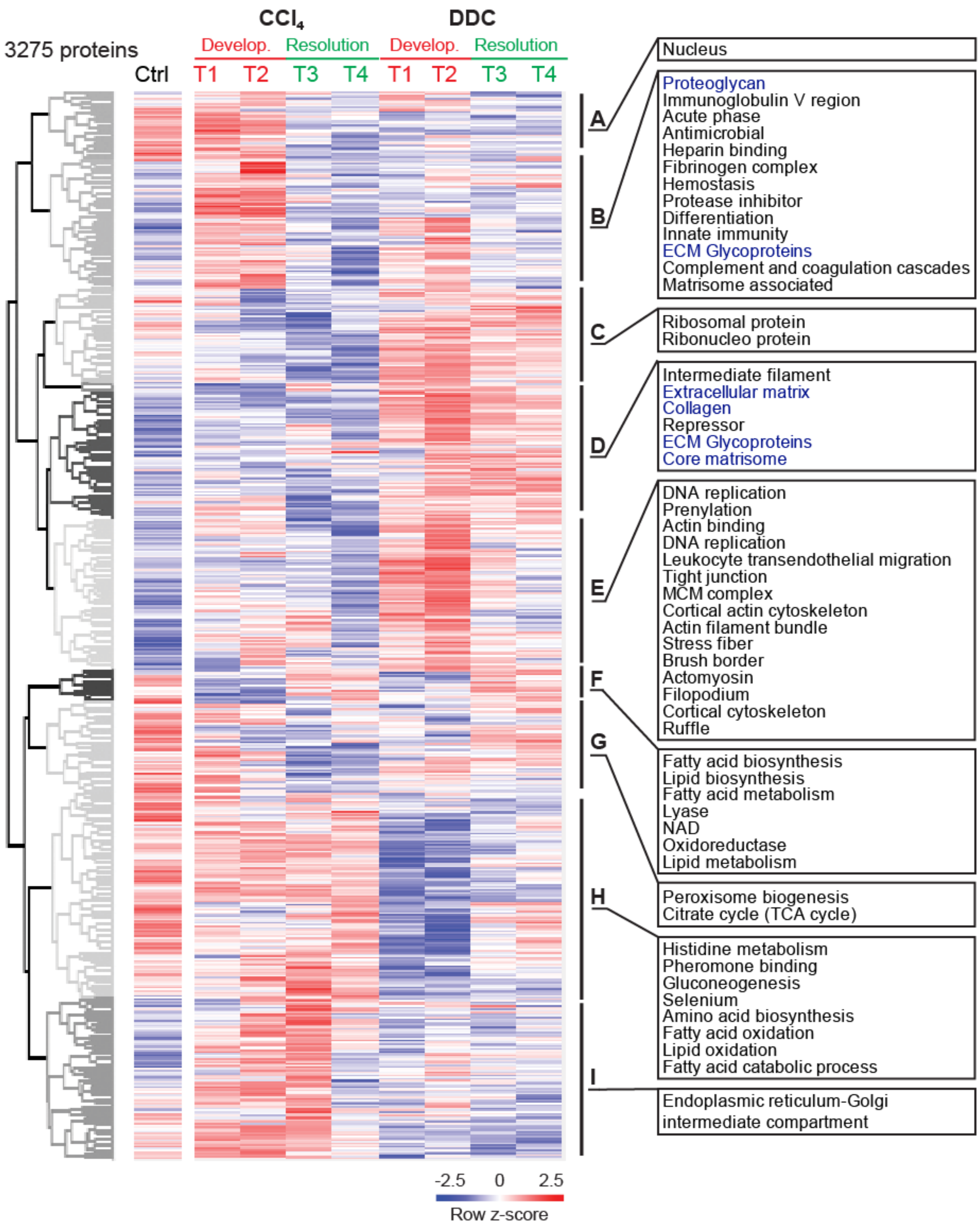


**FIGURE S2** Hierarchical cluster analysis shows the separation of $CCl_4$- and DDC-derived Total proteomes. Unsupervised hierarchical clustering of z-scored mass spectrometry intensities (log 2) of proteins from Total $CCl_4$ and DDC proteomes. UniProtKB keyword enrichment annotations for clusters of proteins (Benjamini-Hochberg FDR < 0.04) are shown (ECM-related keywords in blue).

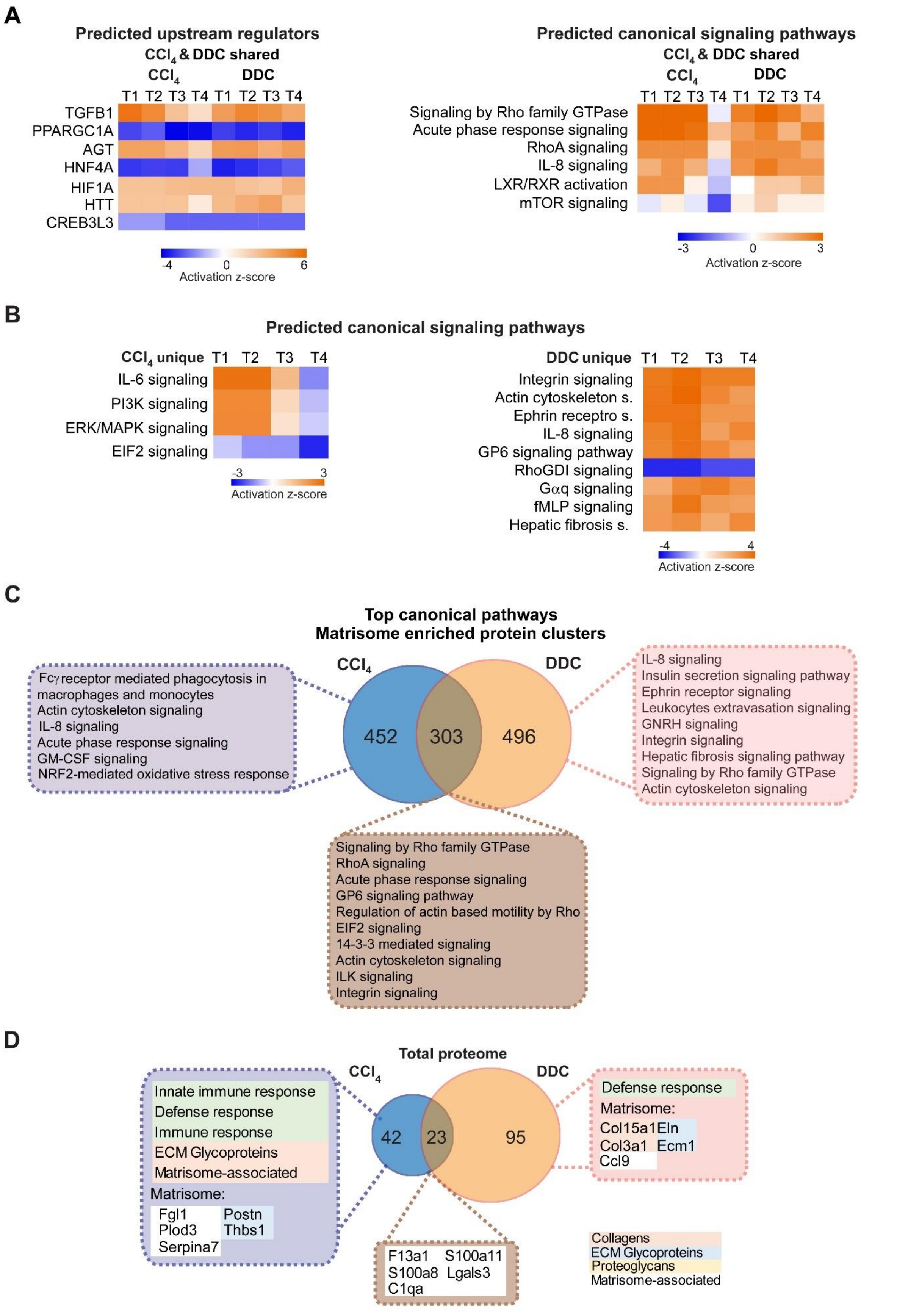


**FIGURE S3** Ingenuity Pathway Analysis (IPA) of upstream regulators and canonical signaling pathways in $CCl_4$ and DDC models. (**A**) Hierarchical cluster analysis of the activity score of the upstream regulators and downstream canonical signaling pathways at the indicated time points predicted by IPA from 762 proteins shared between $CCl_4$ and DDC protein signatures shown in Figure 2A. (**B**) Hierarchical cluster analysis of the activity score of the canonical signaling pathways at the indicated time points predicted by IPA from proteins uniquely identified in $CCl_4$ (514) and DDC (1074) Total proteomes. (**C**) Venn diagram compares proteins from matrisome-annotated clusters shown Figure 2A. Top canonical signaling pathways identified by IPA are shown. (D) Venn diagram compares proteins identified as newly induced by fibrosis development in $CCl_4$ and DDC models, highlighting thus qualitative changes in total proteomes induced by fibrogenesis. We identified 65 newly induced proteins in $CCl_4$- and 118 in DDC-treated livers. Top canonical signaling pathways identified by IPA associated with $CCl_4$-induced proteins and selected matrisome annotated proteins are shown. Matrisome proteins are highlighted with color coding to indicate identified matrisome categories.

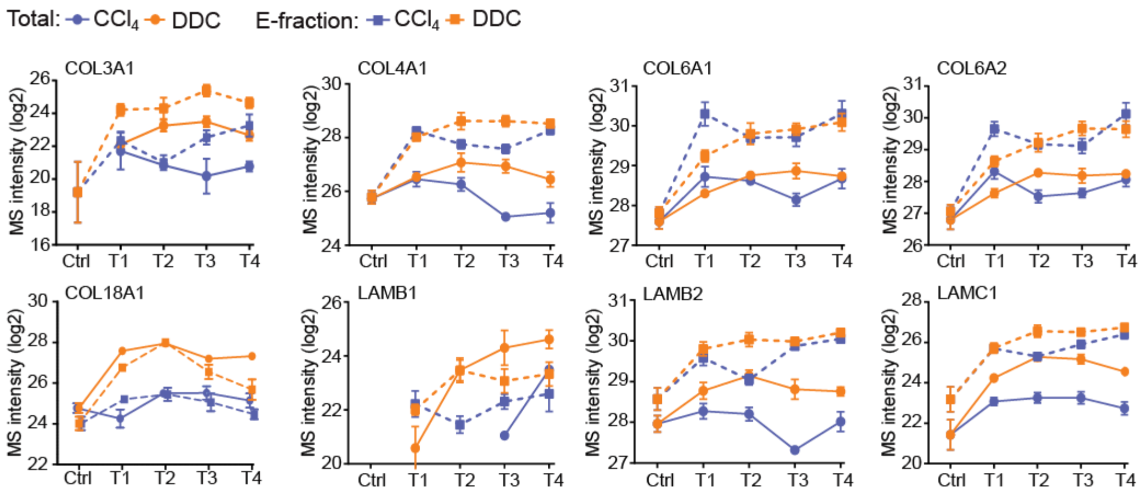


**FIGURE S4** Collagens and laminins identified in Total proteomes. The line plots show the time-dependent change in mass spectrometry intensities of indicated selected collagens and laminins in Total (solid line) and E-fraction (broken line) proteomes in $CCl_4$ and DDC models; n = 4–6.

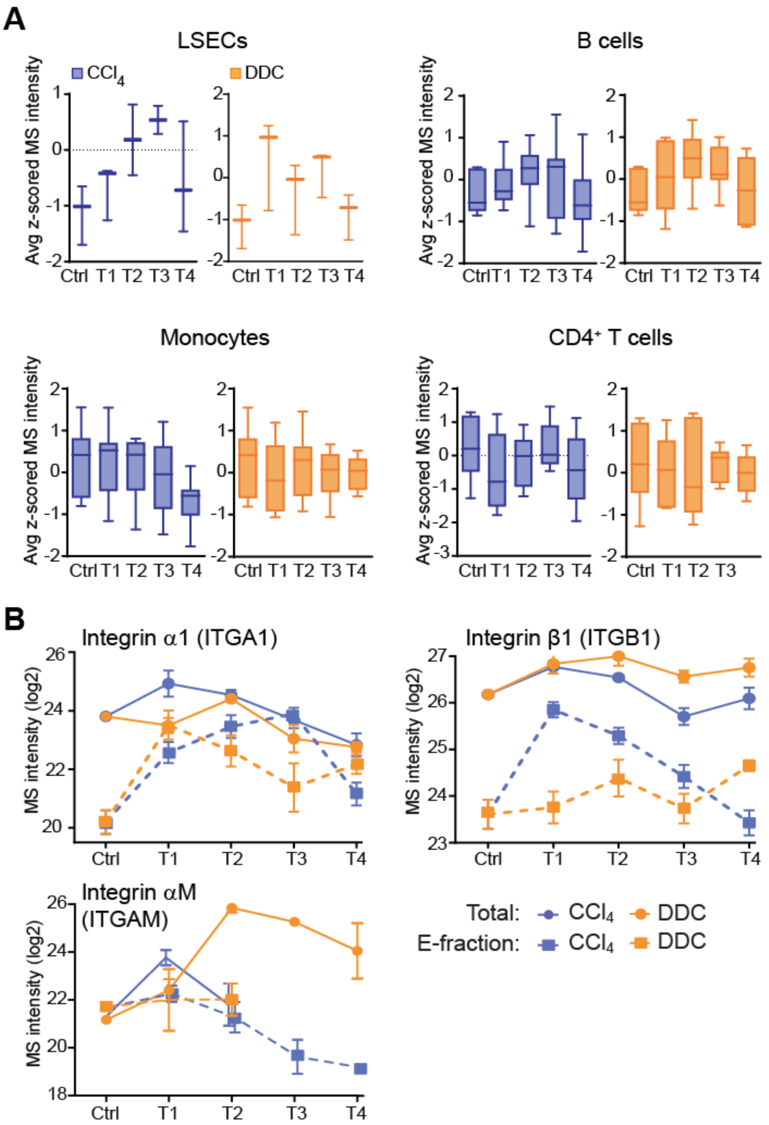


**FIGURE S5** Dynamics of LSECs and immune cells with their cell type-specific integrin expression profiles. (**A**) Box plots show z-scored mass spectrometry (MS) intensities of the indicated cell type-specific protein signatures and time. LSECs (n = 3), B cells (n = 7), monocytes (n = 12), and CD4⁺ T-cells (n = 6). (**B**) The line plots show the change in MS intensities of indicated selected integrins in Total (solid line) and E-fraction (broken line) proteomes in $CCl_4$ and DDC models; n = 4–6.

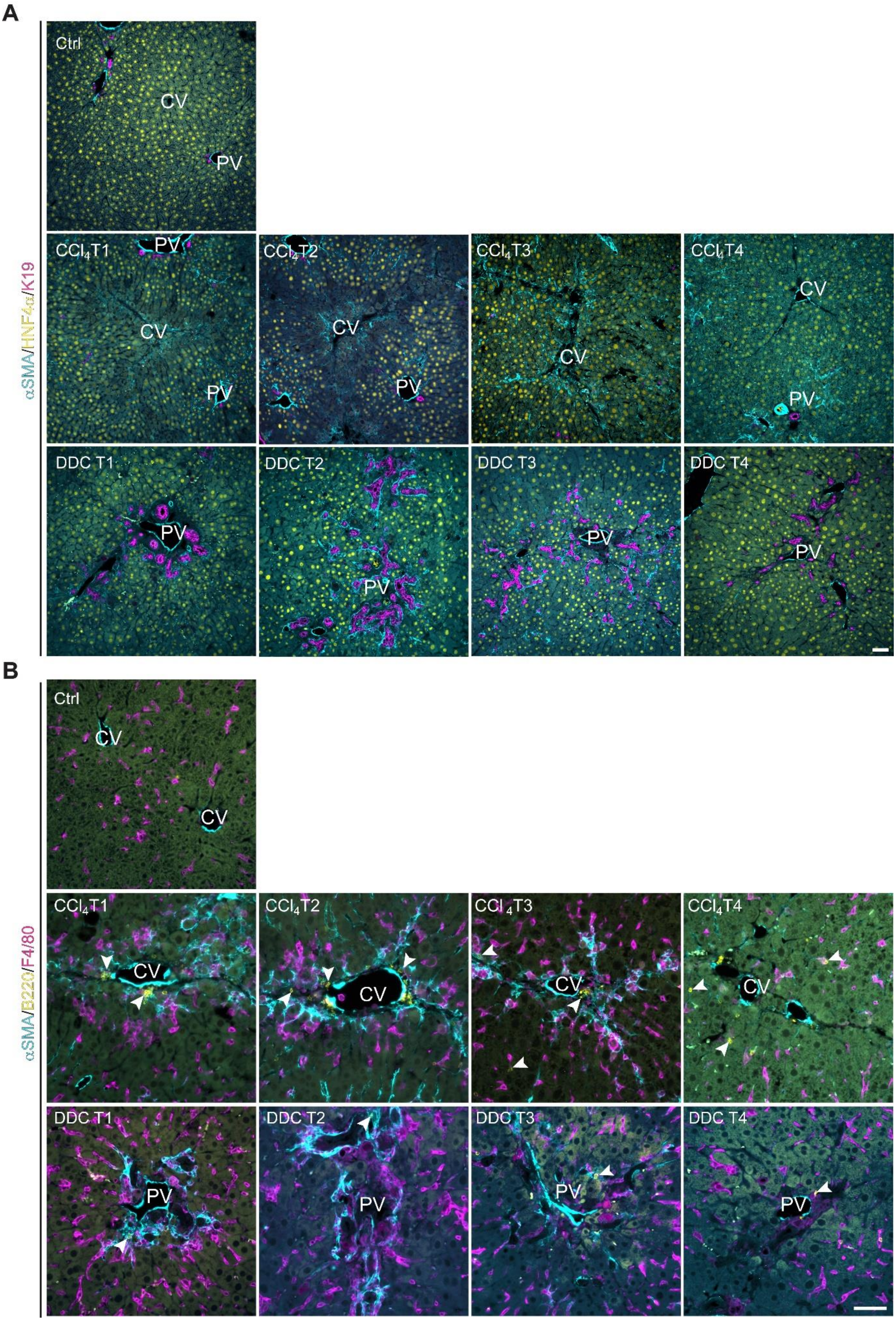


**Figure S6.** Liver cell-type specific dynamics development during fibrogenesis and healing documented by immunofluorescence microscopy corroborates the MS-derived cell-type specific signature dynamics. (**A-B**) Representative IF images of liver sections from untreated controls (Ctrl), $CCl_4$-, and DDC-treated mice at time points of fibrosis development (T1, T2) and resolution (T3, T4) immunolabeled for indicated cell-type specific markers. (**A**) HNF4$\alpha$ (yellow; detecting changes in hepatocyte abundance), $\alpha$SMA (cyan; detecting activated HSCs and portal fibroblasts), and CK19 (magenta; detecting cholangiocytes). (**B**) B cell marker (B220, yellow) and macrophage/monocyte/Kupffer cell marker (F4/80; magenta) together with $\alpha$SMA (cyan) antibody show the role of the inflammation component in the context of fibrosis development and resolution in both models. Arrowheads, B cells recruited to the sites of injury. CV, central vein; PV, portal vein. Scale bar = 50 $\mu$m.

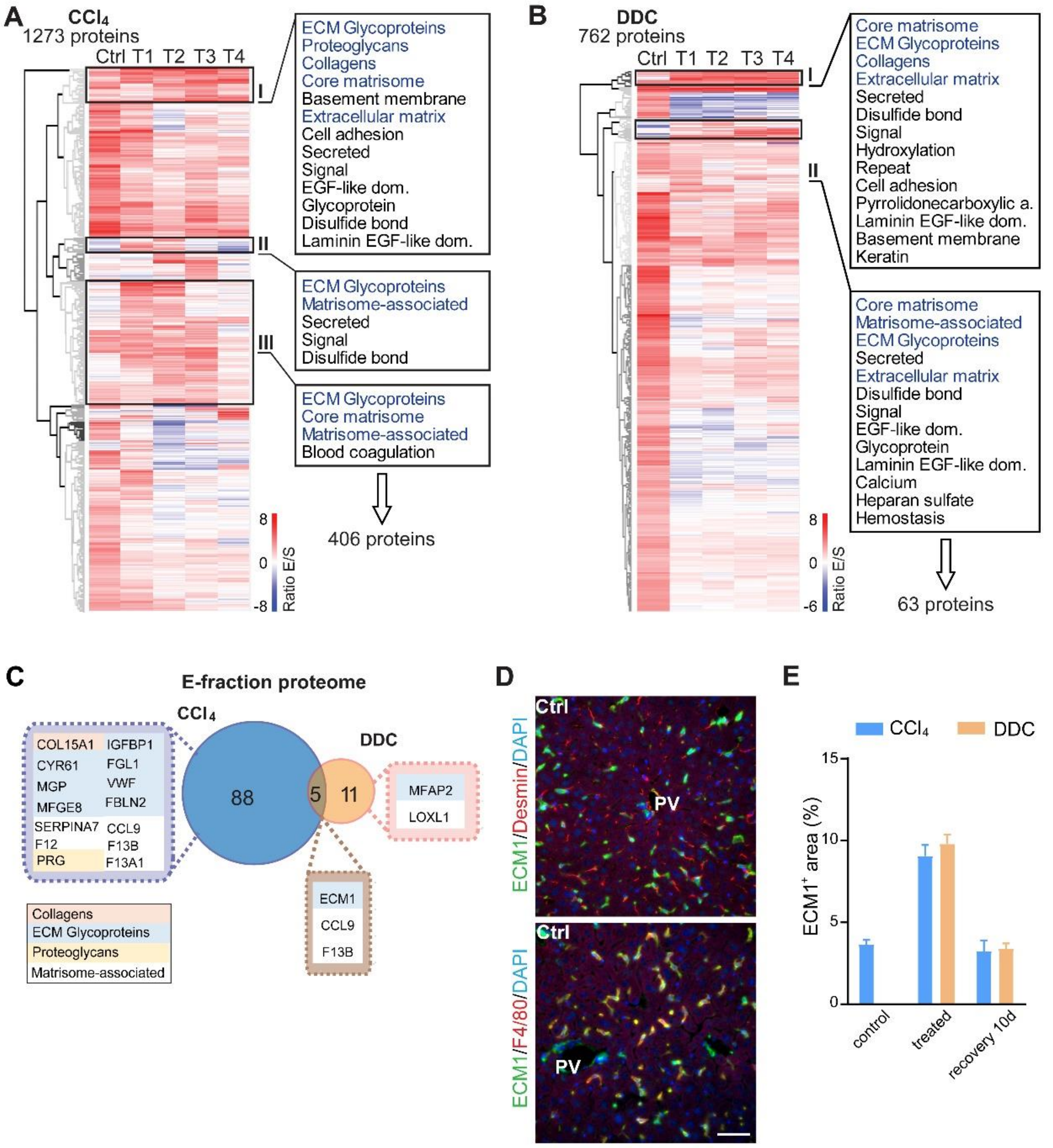


**FIGURE S7** Analysis of matrisome proteins identified in $CCl_4$- and DDC-derived E-fraction proteomes. (**A,B**) Unsupervised hierarchical clustering of mass spectrometry intensity ratios (E-fraction/S-fraction) of proteins differentially expressed in E-fraction in $CCl_4$ (A) and DDC (B) model. UniProtKB keyword enrichment annotations for clusters of proteins ($CCl_4$ clusters I.-III. in A; DDC clusters I. and II. in B) with decreasing solubility over the course of fibrosis are shown (ECM-related keywords in blue). (**C**) Venn diagram shows a comparison of matrisome proteins detected during fibrosis development (T1 and T2) but not in control samples in E-fraction proteomes of $CCl_4$ and DDC model. Color coding indicates identified matrisome categories. (**D**) Representative immunofluorescence images of liver sections from untreated controls (Ctrl) immunolabeled for ECM1 (green) and desmin (red; left panel) or F4/80 (red; right panel). Nuclei were stained with DAPI (blue). Note the colocalization of ECM1 with F4/80 positive cells (in yellow; right panel). PV, portal vein. Scale bar = 50 μm. (**E**) Bar graph shows the quantification of percentage of $ECM1^+$ area per field of view (using 40x objective) within the injury affected area (CV area in $CCl_4$ model and PV area in DDC model) at selected time points.

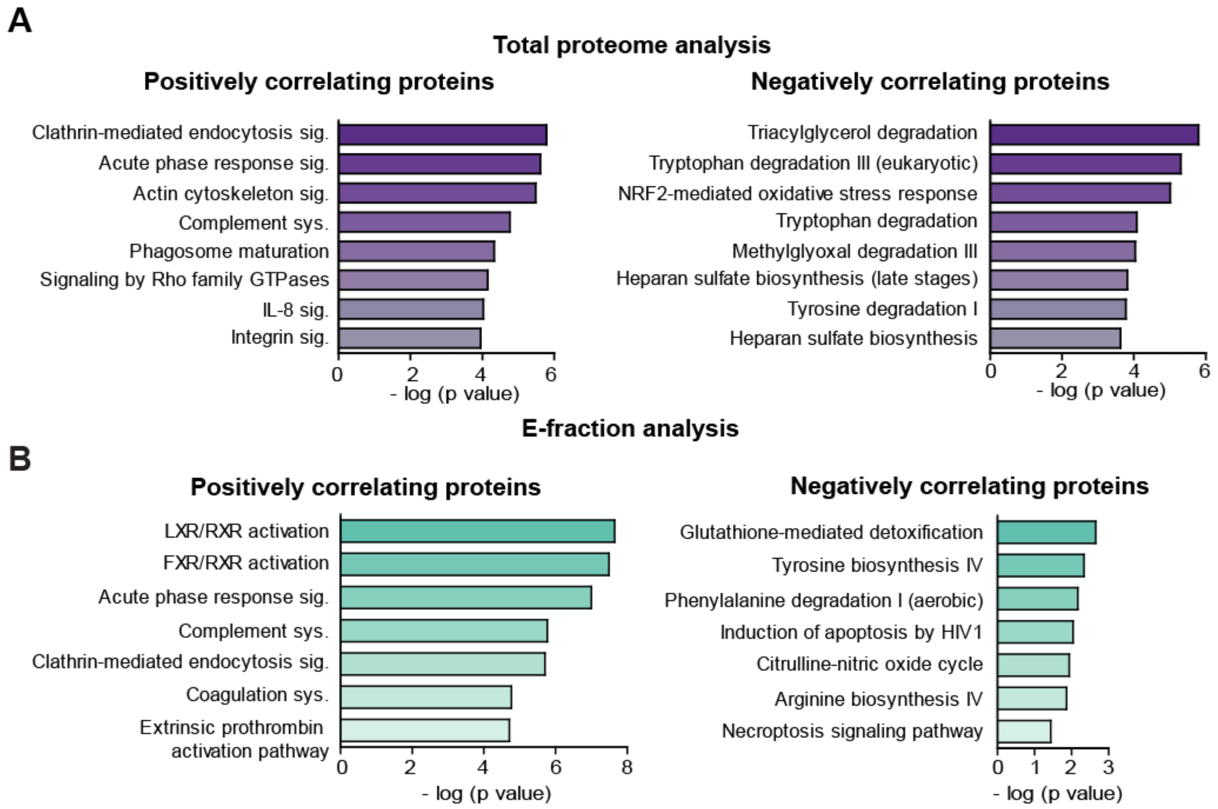


**FIGURE S8** Ingenuity Pathway Analysis (IPA) canonical signaling pathways predicted from correlation analysis. (**A,B**) Canonical signaling pathways identified (IPA) for proteins with significant correlation with fibrous ECM deposits from $CCl_4$-derived Total (A) and E-fraction (B) proteomes.

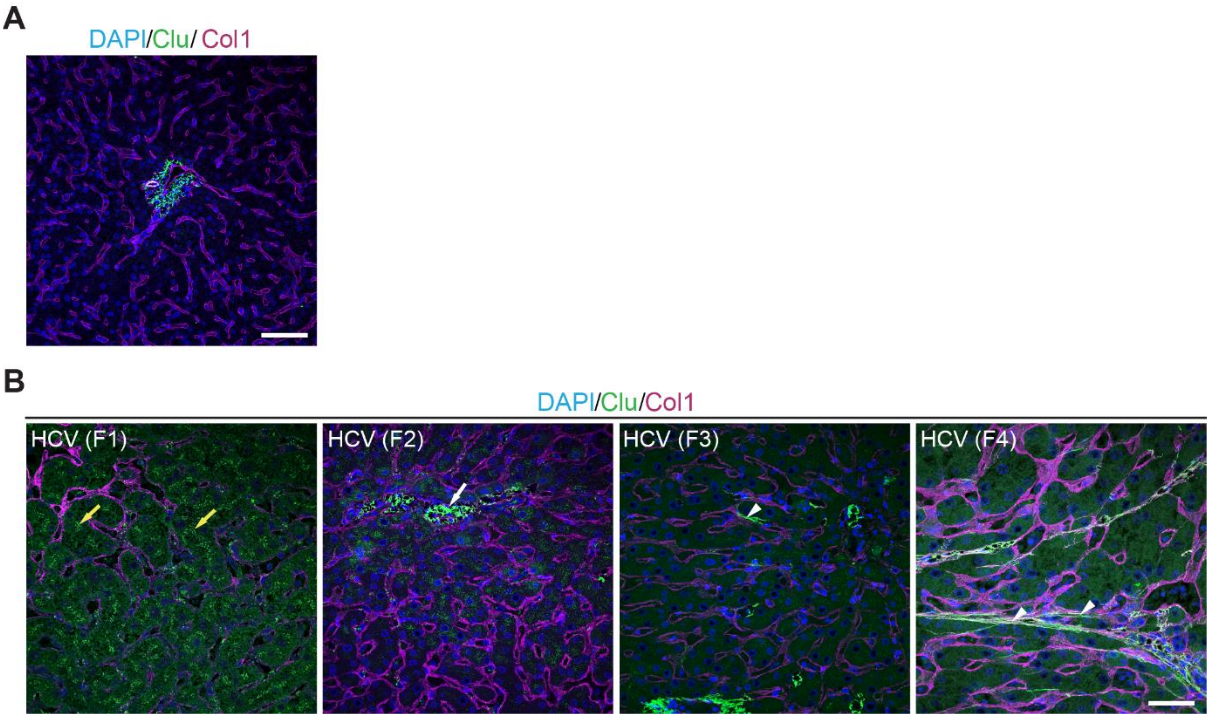


**FIGURE S9** Clusterin expression in control and diseased human livers. (**A**) Representative immunofluorescence (IF) images of control human liver sections immunolabeled for clusterin (green) and collagen 1 (Col1, magenta). Nuclei were stained with DAPI (blue). Note no clusterin expression in healthy parenchyma. Scale bar = 50 μm. (**B**) Representative IF images of human liver sections from different stages of chronic HCV infection immunolabeled for clusterin (green) and collagen 1 (Col1, magenta). Nuclei were stained with DAPI (blue). The images document change in clusterin staining pattern with fibrosis progression from stage F1 to stage F4 according the METAVIR grading system (F1, portal fibrosis; F2, periportal fibrosis; F3, bridging septal fibrosis; F4, cirrhosis). Note clusterin positive bile canaliculi in stage F1 (yellow arrows); clusterin-positive granuli associated with sinusoids in F2 (arrow); clusterin positive collagen fibers within sinusoids in F3 (arrowhead); clusterin positive collagen fibers within parenchyma in F4 (arrowheads). Scale bar = 25 μm.

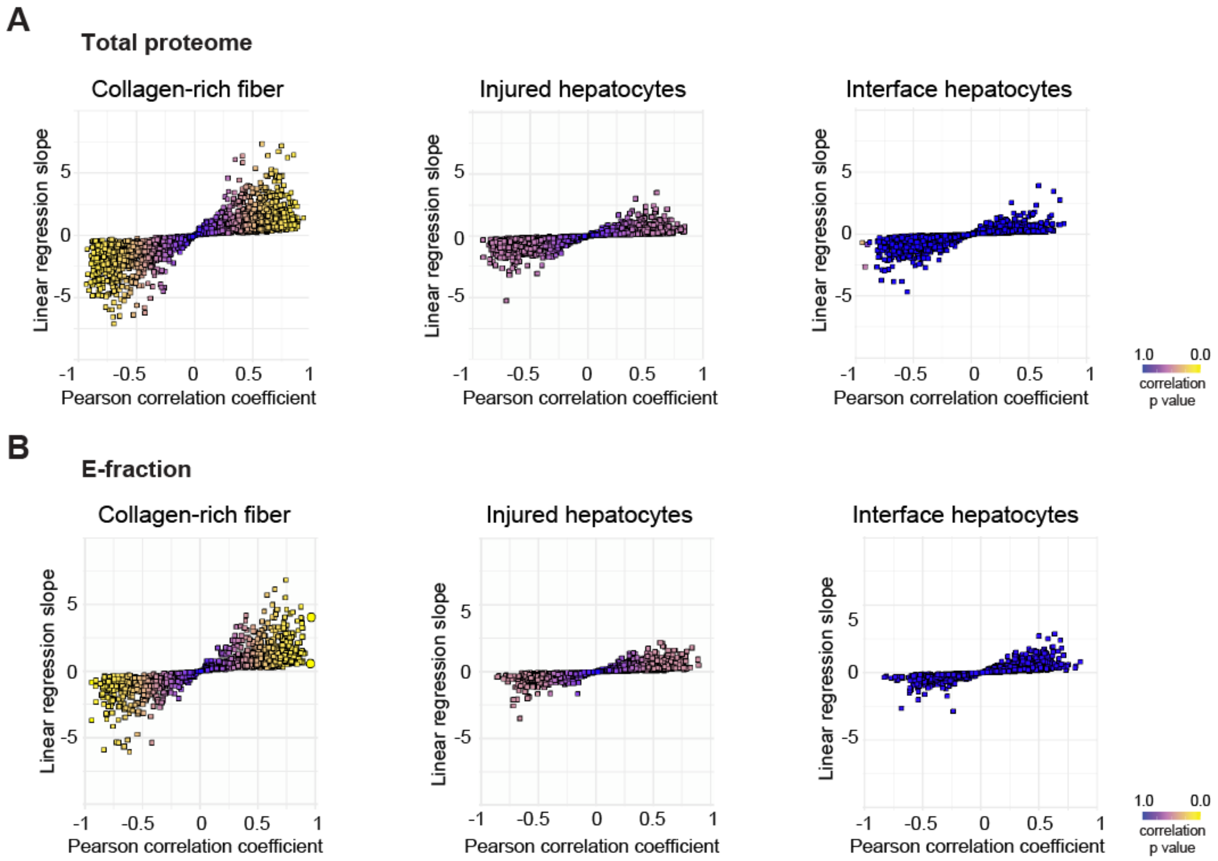


**FIGURE S10** Correlation analysis of protein abundance change and changes in mechanical properties of the local mechanics of liver tissue upon hepatotoxic injury. (**A,B**) The scatter plots show the linear regression slope and the Pearson correlation coefficient for proteins of $CCl_4$ Total proteome (A), and $CCl_4$ E-fraction proteome (B) with average Young's moduli of collagen-rich areas, injured hepatocyte areas, and interface hepatocyte areas. The statistical significance of the correlation is color-coded as indicated.

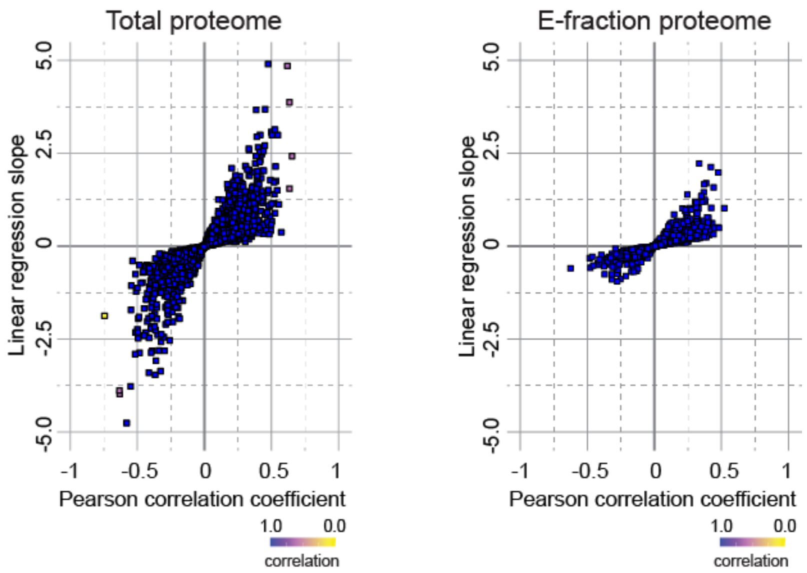


**FIGURE S11** Correlation analysis of protein abundance and changes in fibrotic deposits in the cholestatic model. The scatter plots show the linear regression slope and the Pearson correlation coefficient for proteins of DDC Total (left panel), and E-fraction (right panel) proteomes. The statistical significance of the correlation is color-coded as indicated.

**5. Supporting TABLES**

**TABLE S1. Median Young's moduli for each defined area and treatment.**

| | Median Young's Modulus (kPa) | | |
|---|---|---|---|
| Treatment | Collagen Area | Affected Hepatocyte Area | Interface Hepatocyte Area |
| Control (substitute area) | 1.6 | 1.3 | 1.2 |
| 3 week $CCl_4$ | 2.7** | 1.9** | 0.8** |
| 6 week $CCl_4$ | 4.4** | 2.8** | 2.0* |
| 6 weeks + 10 days | 3.0 | 2.5* | 1.4 |

Statistical analysis was performed on frequency distribution data using Kruskal-Wallis test followed by Dunn's multiple comparison post-testing. ** $p < 0.01$ vs. control; * $p < 0.05$ vs. control.

**Table S2. Antibodies used in the study**.

| Target | clone | supplier | cat number | Reference; RRID# |
|---|---|---|---|---|
| K19 | TROMA III | DSHB | TROMA III | 6933460; RRID:AB_2133570 |
| αSMA | 1A4 | DAKO | M0851 | RRID:AB_2223500 |
| ECM1 | F-1 | Santa Cruz Biotechnology | sc-365335 | RRID:AB_10847810 |
| clusterin | | R&D Systems | AF2747 | RRID:AB_2083314 |
| clusterin | A-11 | Santa Cruz Biotechnology | sc-166831 | RRID:AB_2245186 |
| fibronectin | | Abcam | ab2413 | RRID:AB_2262874 |
| collagen I | | Abcam | ab21286 | RRID:AB_446161 |
| collagen IV | | BioRad | 2150-1470 | |
| integrin αV | EPR16800 | Abcam | ab179475 | RRID:AB_2716738 |
| B220 | RA3-6B2 | BioLegend | 103203 | |
| F4/80 | D2S9R | Cell Signaling | 70076 | RRID:AB_2799771 |
| desmin | | ThermoFisher Scientific | PA5-16705 | RRID:AB_10977258 |
| Alexa Fluor® 488 AffiniPure F(ab')$_2$ Fragment Donkey Anti-Goat IgG (H+L) | | Jackson ImmunoResearch | 705-546-147 | RRID:AB_2340430 |
| Alexa Fluor® 594 AffiniPure F(ab')$_2$ Fragment Donkey Anti-Rat IgG (H+L) | | Jackson ImmunoResearch | 712-586-150 | RRID:AB_2340690 |
| Alexa Fluor® 647 AffiniPure Donkey Anti-Rabbit IgG (H+L) | | Jackson ImmunoResearch | 711-605-152 | RRID:AB_2492288 |
| Alexa Fluor® 488 AffiniPure Donkey Anti-Mouse IgG (H+L) | | Jackson ImmunoResearch | 715-545-150 | RRID:AB_2340846 |
| Alexa Fluor® 488 AffiniPure Donkey Anti-Rabbit IgG (H+L) | | Jackson ImmunoResearch | 711-545-152 | RRID:AB_2313584 |
| Rhodamine Red™-X (RRX) AffiniPure Donkey Anti-Mouse IgG (H+L) | | Jackson ImmunoResearch | 715-295-151 | RRID:AB_2340832 |
